# ATF4 orchestrates IL-1α-induced senescence in adult neural stem cells

**DOI:** 10.1101/2024.03.17.585394

**Authors:** Susanne Neumann, Eric P. Thelin, Sreenivasa Raghavan Sankavaram, Sanna Persson, Leonor Fonseca, Noah Moruzzi, Ellen Iacobaeus, Maria Bergsland, Elena Popova, Michael Bader, Mikael Svensson, Alexander Espinosa, Ruxandra Covacu, Lou Brundin

**Affiliations:** Department of Clinical Neuroscience, Center for Molecular Medicine, Karolinska University Hospital, Karolinska Institute, Stockholm, Sweden; Department of Clinical Neuroscience, Karolinska Institute, Medical Unit Neurology, Karolinska University Hospital, Stockholm, Sweden; The Rolf Luft Research Center for Diabetes and Endocrinology, Karolinska Institutet, Stockholm, Sweden; Department of Cell and Molecular Biology, Karolinska Institute, Stockholm, Sweden; Max-Delbrück-Center for Molecular Medicine in the Helmholtz Association (MDC), Berlin, Germany; Charité University Medicine, Berlin, Germany; German Center for Cardiovascular Research (DZHK), Partner Site Berlin, Germany; Institute for Biology, University of Lübeck, Lübeck, Germany; Department of Clinical Neuroscience, Neurosurgery Unit, Karolinska University Hospital, Karolinska Institute, Stockholm, Sweden; Department of Medicine, Center for Molecular Medicine, Karolinska University Hospital, Karolinska Institute, Stockholm, Sweden

## Abstract

Adult neural stem cells (NSC) are a potential source for the regeneration of damaged tissue during neuropathological conditions, but much remains unexplored. In an attempt to study the influence of neuroinflammation on NSCs, we generated a transgenic reporter rat strain that expresses the *Discosoma sp*. red (DsRed) fluorophore in NSCs and subjected it to traumatic brain injury (TBI). Transcriptomic analysis of NSCs isolated from TBI revealed an enrichment of stress response genes that pertained to endoplasmic reticulum (ER) stress and integrated stress response (ISR). Downstream analysis on NSC cultures pinpointed IL-1α as a trigger of ISR in these cells. At concentration levels similar to the ones measured post-TBI in rats, IL-1α induced the translation of activating transcription factor 4 (ATF4), an ISR master regulator. Further, ATF4 was necessary for the IL-1α -dependent induction of a senescent profile in NSCs, which included a metabolic shift towards glycolysis, induction of senescence-associated secretory phenotype, SASP, and cell cycle arrest. In summary, the ISR/ATF4 pathway seems to play a major role in NSC function during neuroinflammation and provides a therapeutic tool for protecting the NSC pool during these conditions.

## Introduction

Neural stem cells (NSCs) in the adult central nervous system (CNS) are capable of self-renewal and generation of mature neurons, oligodendrocytes and astrocytes. NSCs are actively dividing in the subventricular zone (SVZ) [1] and the subgranular zone (SGZ)[2] in the brain and are found in a quiescent state in the spinal cord [3]. These NSC niches are under strict regulatory cues ensuring a balance between cell proliferation and quiescence, thus providing sufficient numbers of cells for differentiation while preserving the NSC pool. The regenerative potential of NSCs has been demonstrated in numerous studies on traumatic and autoimmune disease models [4]. NSCs migrate to the injury site and give rise to mature cell types that could potentially replace the lost or damaged tissue or contribute with trophic support [5]. Unfortunately, these pathological events have long-term consequences on NSC function [6, 7]. Understanding how NSCs are affected by neuroinflammatory conditions, such as traumatic brain injury (TBI) and multiple sclerosis (MS), is key to finding therapeutical options that will protect the niche from NSC depletion and maintain or even improve their performance during these pathological conditions.

Cellular stress pathways, in particular the endoplasmic reticulum (ER) stress and integrated stress response (ISR), have gained increased attention for their involvement in several neuropathological diseases including Alzheimer’s (AD) and Parkinson’s disease (PD)[8], multiple sclerosis (MS) and traumatic brain injury (TBI) [9]. The ISR is triggered by various stimuli such as, viral infections, misfolded proteins and amino acid deprivation leading to the activation of four key enzymes that then converge into the activation of eukaryotic translation initiation factor 2a (eIF2α) and the translation of activating transcription factor 4 (ATF4).

ATF4, in turn, induces expression of genes necessary for the alleviation of transient stress or leads to induction of apoptosis in case chronic stress occurs. ATF4 dimerizes with other transcription factors, which will influence the downstream outcome of its activation. This explains the conflicting reports on the outcome of ISR/ATF4 involvement during neuropathological conditions. Both beneficial effects, such exerting a protective effect on oligodendrocytes in MS and its animal model [10] or detrimental effects, as being associated to AD [11] have been attributed to the ATF4 involvement. In NSC, the ATF4 engagement has been mainly studied during homeostatic conditions in embryonic NSCs where it seems to influence cell cycle regulation and neurogenesis [12]. The role of ATF4 in NSC during inflammation is understudied.

In the current work, we reveal the activation of ISR in NSCs during TBI. Further, we pinpoint the IL-1α as a trigger for this pathway. This was achieved by using a new transgenic reporter rat with *Discosoma sp*. red fluorophore (DsRed)-expressing brain NSCs, which enabled their isolation and analysis *ex vivo*. Together with *in vitro* NSC culture experiments, and using transcriptomics, functional perturbations, and cell function analysis, we describe the induction of a senescent phenotype in adult NSCs triggered by IL-1α through ISR/ATF4 activation.

## Material and methods

### Animals

This study was conducted with the approval by the ethical committee of the Swedish Board of Agriculture. Animals were housed in a 12h light/dark cycle with food and water *ad libitum*.

### Generation of the RedSox2 transgenic reporter rat

The RedSox2 rat strain was generated by using standard cloning techniques. Firstly, to identify the rat sox2 promoter region, we aligned the mouse 5.5kb promoter region and enhancer sequences previously described in mouse [13–16] to the rat *Sox2* sequence region upstream of the *Sox2* transcription start site (TSS) (rat genome assembly rn4, Baylor College of Medicine HGSC v 3.4). A 6kb genomic region upstreams from the *Sox2* TSS was identified as the best corresponding sequence to the mouse 5.5 kb which contains the *Sox2* Regulatory Region 1 (SRR1). To verify the functional relevance of this rat sequence region, transcription factor binding sites (TFBS) were identified using ConSite (http://consite.genereg.net/cgi-bin/consite) confirming similar TFBS as in 5.5kb region mouse. The rat 6kb region was amplified from the BAC clone CH230-350H23 (http://bacpac.chori.org), subcloned into TOPO-XL (Invitrogen) and then further subcloned into pDsRed-Express2 (Clontech, cat no 632535). Due to the GC-rich region being close to the *Sox2* TSS, the 6kb region was amplified in two portions (1-3669 bp and 3670-6000bp) and then fused together. The second distal enhancer (SRR2) was cloned into the pdDsRed-Express2, (Figure 1).

**Figure 1.**
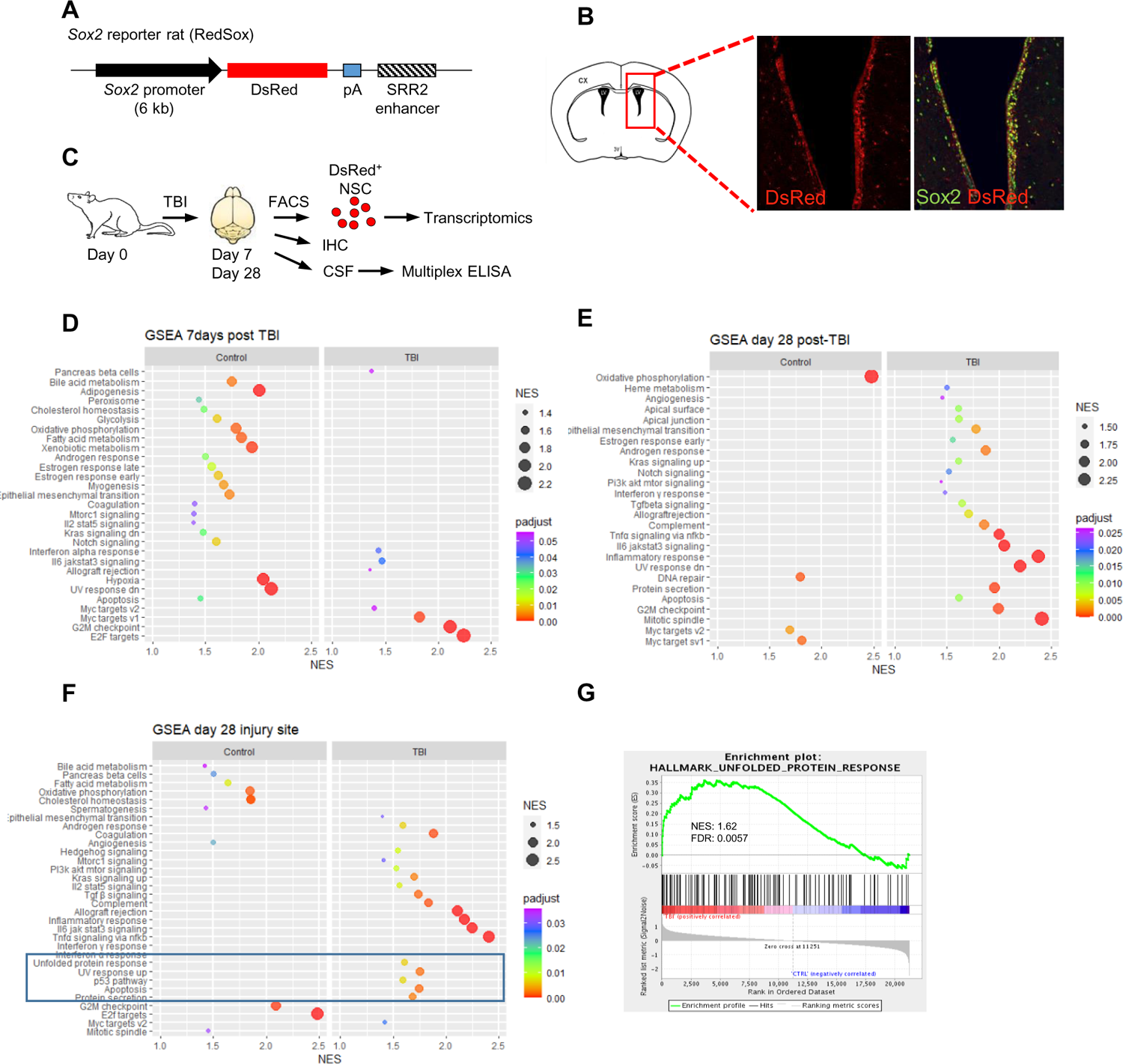
(**A**). Schematic view on DNA construct used to generate the reporter rat. A 6kb sequence upstream of the TSS + of the Sox2 gene was cloned in front of the DsRed-Express-2 fluorophore sequence. The enhancer sequence of (SRR2) regulating the Sox2 expression was also added to the construct. The construct was then microinjected into the male pro-nucleus of DA/HanRj rats zygotes. (**B**). Schematic depiction of coronal sections of rat brains slides depicting the lateral ventricles, one marked with a red border. Histological sections of the RedSox reporter rat showing an over-view of DsRed reporter expression (red cells) in the lateral ventricle. Sox2 immunolabeling (green) overlaps with DsRed expressing cells, confirming the DsRed reporter function. (**C**). Experimental set-up for the TBI. (**D-F**).Bubble plot of GO terms enriched in sorted NSC from TBI or healthy rat brains 7days (**D**), 28 days(E) or 28 days injury site(**F**) post-TBI. The blue rectangle highlights different stress and cell death pathways. (**G**) Enrichment plot showing enrichment of UPR genes in TBI-isolated NSCs. Padjust=adjusted pvalue; NES= normalized enrichment score; FDR=false discovery rate.

Construct microinjection into the pronucleus of Dark Agouti (DA)/HanRj rat zygotes was performed at the Max-Delbrück Center for Molecular Medicine using established technologies [17]. Efficiency of transgenic rat production is independent of transgene-construct and overnight embryo culture. The offspring was genotyped by PCR (using the following primers: Fw: CAAGTCAATCTACATGGCCAAGAA, Rev: TTCAAAGTTTCTT TTATTCCTATG) and a positive founder was mated to establish the transgenic RedSox2 line.

### Traumatic Brain Injury (TBI)

The controlled cortical impact (CCI) model, an experimental TBI that mimics a human focal brain injury [18], has been described in detailed elsewhere [19]. In short, the animals were anesthetized using a mixture of 5% isoflurane, intubated, and mechanically ventilated with a maintenance dose of 2–3% isoflurane, monitored using a pulsoximetry device (MouseSTAT™, Kent Scientific, Torrington, CT, USA). Following adequate sedation, local anesthesia [0.15 mL of bupivacaine (Marcaine®) 0.25%] was injected subcutaneously in the cranial midline, while buprenorphine (Temgesic®) (0.05 mg/kg) and carprofen (Rimadyl®) (5 mg/kg) were injected subcutaneously in the abdominal region to provide analgesia. Eye-gel [containing fusidic acid (Fucithalmic®)] was used, and isotonic saline (NaCl 9 mg/mL) was continuously used to rinse the wound. The animal was placed on a heating pad attached to a stereotaxic frame (Model 900, Agnthos, Stockholm, Sweden). The body temperature was maintained at around 37°C. A 0.5 mm diameter diamond tip drill (Microspeed 317 IN; Silfradent, Forli, Italy) was used to remove a portion of the parietal bone over the right hemisphere, at about 3.5 mm right of the central suture and 4.5 mm posterior to lambda, using a surgical microscope (Wild Heerbrugg M3C Stereozoom Microscope, Leica, Wetzlar, Germany). A commercially available CCI device (TBI 0310, Precision Systems and Instrumentation LLC, Lexington, KY, USA) with a 3 mm in diameter piston was used to impact a 3-mm deep lesion in the right parietal lobe (a lesion defined as a “severe TBI” [REF: 27604719]). The wound was sutured using Vicryl 4-0 (Ethicon, Johnson & Johnson, New Brunswick, NJ, USA) and the animals were placed on a heat pad until fully awake (∼30 min) before returning to their home cages.

### Florescence-Activated Cell Sorting (FACS)

#### Ex vivo FACsorting of DsRed^pos^ NSCs

Following TBI, the animals were kept for 7 or 28 days after which they were euthanized using CO2. The brains were extracted and the SVZ were isolated from the ipsi-lateral and contra-lateral hemispheres under a dissection microscope.

The injury area together with 1mm of the surrounding area was also collected from the 28days group. The biopsies were dissociated mechanically using pipette trituration and enzymatic dissociation using 10U/ml papain (Worthington) and DNase (Sigma-Aldrich) as previously described (Covacu et al 2006). The cells were transferred to FACS medium containing PBS/1% bovine serum albumin (BSA)/DNAse and analysed with a BD Influx^TM^ cell sorter. The sorted cells were pelleted at 220Xg for 5min and lysed with Trizol (Invitrogen) for subsequent RNA extraction.

#### Sorting of DsRed^pos^ /DsRed^neg^ from in vitro cultured NSCs

Following their first or second culture passage, the NSCs were sorted into *DsRed^pos^* and *DsRed^neg^* populations by using a BD Influx^TM^ cell sorter. The single-cell suspension was prepared by dissociating the neurospheres using mechanical and enzymatic dissociation, see previous section [20]. The cells were sorted in FACS buffer containing PBS/1%BSA/DNase. Following the FACSorting, the cells after their first passage placed in propagation medium, see ‘neural stem cell’ section, while cells after the second passage were used in experiments.

### RNA extraction, cDNA preparation

RNA isolation for downstream quantitative RT-PCR analysis was performed according to the manufacturer’s protocol for TRIzol (Invitrogen, # 15596018). Equal amounts of RNA from the different samples were used to prepare cDNA by using the iScript kit (Invitrogen #1708891BUN).

The RNA extraction for the global transcriptome analysis was performed by using a modified TRIzol protocol. The biopsies were mechanically homogenized in 1ml TRIzol (Invitrogen, # 15596018) using TissueLyser LT (Qiagen). If cell cultures were used, these were lysed with TRIzol dicrectly in cell culture wells. The upper phase was collected, mixed with isopropanol and glycogen (GlycoBlue coprecipitant, ThermoFisher Scientific #AM9516, end-concentration 50μg/ml) and incubated at −20⁰C overnight. The RNA isolation was finished the next day according to the manufacturer’s protocol. The RNA was further purified and concentrated by using RNeasy MinElute Cleanup Kit (Qiagen #74204). The RNA integrity was analysed with a bio-analyser.

### Affymetrix transcriptome analysis

Gene expression was measured in the following experimental groups: FACSorted NSCs (*DsRed^pos^*) from the SVZ of healthy control animals and TBI-injured animals day 7 and day 28 post-injury, (ipsilateral side) and also from the injury area from the day 28 animals, n=3 in each experimental group and time-point, 18 animals in total. The array platform used was Affymetrix GeneChip RAT Gene ST 2.0. Array hybridization and basic data processing were performed at the Bioinformatics and Expression Analysis Core facility at Karolinska Institutet, Stockholm, Sweden. Data processing involved signal correction of background using the GC composition-based background correction algorithm, array normalization with global median and signal summarization using the probe logarithmic intensity error estimation (plier). GeneChip Expression console from Affymetrix was used. After filtration of the low-expression genes (<50) and non-annotated genes, the data was analysed using Gene Set Enrichment Analysis (GSEA) [21, 22]. The data sets in this manuscript will be deposited on NCBI’s Gene Expression Omnibus.

### RNAsequencing

RNA sequencing was performed on RNA isolated from NSCs with perturbed expression of ATF4 using siRNA (siATF4). The control cultures were transfected with or a non-targeting control siRNA (siNTC). The cultures were stimulated with IL-1α or kept unstimulated and harvested at 4h and 24h post-stimulation. For RNA isolation, see the “RNA extraction” section. RNA quality control, library preparation, sequencing, sequence annotation and analysis was done at the Bioinformatics and Expression Analysis Core facility at Karolinska Institutet. Purified RNA (see RNA extraction) and quality controlled was uses for generating the sequencing libraries using the Illumina TruSeq Stranded mRNA with polyA selection. cDNA libraries were sequenced on a Illumina Nextseq 2000. FASTQ files, generated with CASAVA software were aligned using STAR. Differential gene expression was analyzed with the R package DESeq2. Heatmaps were generated in Multiple Expression Viewer (MeV) [23].The functional analysis and canonical pathway analysis was generated with Ingenuity Pathway Analysis (IPA, Qiagen, http://www.qiagen.com/Ingenuity) and WEB-based Gene Set Analysis Toolkit (WEBGESTALT) [24–27]. Genes from the dataset that met the signal intensity cutoff of >50 and passed a 5% FDR level were considered for the analysis. For the gene set enrichment the fGene Set Enrichment Analysis (GSEA) script for R was used.

### Tissue perfusion and preparation

The rats were placed under deep anesthesia though intraperitoneal injection of a lethal dose of pentobarbital or via inhalation with isoflurane. The animal ware transcardially perfused with 1X PBS and 4% formalin solution. The brain was extracted and post-fixed in 4% formalin solution for 24hrs at 4°C, washed with PBS and placed in 30% sucrose over-night at 4°C. The brains were frozen and coronal sections (14μm) were made in a using a cryostat (Leica, SM2000R).

### Immunohisto/cytochemistry

### Immunocytochemistry

Cells were cultured on PDL (10-20 ug/ml, Sigma) Nunc™ Lab-Tek™ Chamber Slide System were used (#177402, Thermo scienfic). Following the experiment, the cells were fixed using 4%PFA in PBS for 10 minutes in RT. The cells were blocked in PBS containing 5% goat serum (Bio-Rad Antibodies) and 0.1% saponin (SigmaAldrich) for 30 min at RT. The antibodies were diluted in a similar solution but containing 1% serum and incubated at 4°C over-night for the primary and at 37°C for 30 minutes for the secondary antibody, respectively. For intranuclear staining (Sox2, Atf4), an initial permeabilization step with 3% TRITON-X at 4°C for 10 minutes, was introduced prior to the blocking step.

Between incubations the cells were washed in PBS/Saponin and finally mounted in MOWIOL (SigmaAldrich). <u>Immunohistochemistry</u>. The immunolabeling of brain sections started with re-hydration in PBS for a few minutes followed by a blocking step and immunolabeled using the same procedure as for cell cultures. For EdU staining, the manufacturer’s instructions were followed (Thermo Scientific). The immunolabelings were analysed using a Revolve Fluo (ECHO) epifluorescence microscope or a confocal microscope (ZEISS).

Immunolabeling quantifications were performed manually and blinded, either by counting the entire well or several objective fields of view (10X or 20X objective) with similar cell density.

### Gel electrophoresis and western blotting

NSCs attached on PDL-coated 12-well plates (Nunc, #150628) were washed once with ice-cold PBS and lysed in 120 μl of CellLytic M™ cell lysis reagent (Sigma-aldrich #C2978) supplemented freshly added Halt™ protease and phosphatase inhibitor cocktails (Thermo Fisher Scientific #78440). The cells were scraped off the plate, sonicated (Sonoplus, Bandelin) and centrifuged at 10,000Xg for 10 minutes to remove cell debris. The protein concentration was measured using Pierce™ 660 protein assay reagent (Thermo fisher scientific #22660). The debris-cleared supernatant was mixed with Laemmli sample buffer 4X (Bio-Rad, #1610747) supplemented with 50mM DTT (Sigma-Aldrich, #10197777001) and heated at 95°C for 5min. Equal quantities of proteins from each individual sample were separated on mini-Protean® TGX™ mini pre-cast gels (BioRad, #4561023) at 120V until the leading dye reached the bottom of the gel. The proteins were transferred to a PVDF transfer membrane (0.45 μm, ThermoScientific #88518) by using a Trans-Blot®Turbo™ transfer system (Bio-Rad) at a constant current of 2.5A for 30 minutes. The membranes were blocked using TBS/0.1% Tween (Sigma-Aldrich) with 5% of bovine serum albumin (Sigma-Aldrich) or nonfat dried milk (Panreac AppliChem). The membrane was incubated overnight at 4°C with the primary antibody diluted according to the manufacturer’s instructions. Following washing with TBS/T the secondary HRP-coupled antibody was applied for 1h at room temperature. The band detection was achieved by using Clarity™ Western ECL substrate or Clarity™ Max Western ECL substrate (Bio-Rad). The band intensities were analysed with Image Lab™ software (Bio-Rad).

### Antibodies

Primary antibodies used for WB: anti-ATF4 (D4B8, 11815S, 1:1000 for WB and 1:200 for immunolebeling) anti-phospho-eIF2α (ser 51, D9G8, #3398) anti-eIF2α (#5324) all diluted 1:1000 and purchased from Cell Signaling; anti-p21 (1:1000, EPR3993, ab109199, Abcam) anti-olig2 (1:2000, EPR2673, ab109186, Abcam), anti-phos-Olig2 (ser10, ser13, ser14; 1:1000 # PA5-35405, Invitrogen). Secondary HRP-linked antibodies: anti-mouse IgG (1:3000, #7076) anti-rabbit IgG (1:3000, #7074) from Cell Signaling. Anti-Beta-Actin conjugated with HRP 1:25,000 (Sigma-Aldrich, A3854 (clone AC-15), anti-Sox2 (Millipore Sigma, AB5603, 1:100).

### Multiplex Elisa

Cerebrospinal fluid (CSF) was collected using a butterfly needle via the cisterna magna of rats deeply sedated with Dormicum® (midazolam) and previous to rat euthanasia using cardiac perfusion with PBS and PFA. CSF was centrifuged cold at 10,000 g for 5 min. The CSF supernatant was stored at −70 degrees. CSF contaminated with blood was discarded. For analysis the Bio-Flex Pro™ Rat Cytokines Assay (Bio-Rad, 24-plex panel #171-K1001M) was used on a Bio-Plex^®^ MAGPRIX™ multiplex reader and analysed with Bio-Plex Analysis software (Bio-Rad).

### NSC culture and cytokine stimulation and differentiation

The NSC cultures were prepared as previously described [20]. The SVZ were extracted from RedSox2 rats (bred in house). After dissociation, the cells were cultured in propagation medium containing: DMEM/F-12 medium (Life technologies cat no 31331-028) with B27 supplement without vitamin A (Life Technologies, cat no 12587-010), 100U/mL Penicillin-Streptomycin (Life Technologies), 20ng/ml epidermal growth factor (mouse EGF, Sigma-Aldrich) and 10ng/ml human basic fibroblast growth factor (bFGF, R&D systems). For the RedSox2 NSC characterization, the neurospheres were triturated into single cell solutions and FACsorted (FACS, BD Influx^TM^ cell sorter) into DsRed^pos^ and DsRed^neg^ populations, either after their first or second culture passage. All NSCs, sorted or not, were used in the experiments after the second passage. The cells were plated onto poly-D-lysine (PDL, Sigma-Aldrich) coated culture plates or glasses. The following steps, including cytokine stimulation and transfection, were performed in propagation medium, unless differently specified. The cytokine stimulation with rat recombinant IL-1α (R&D, # 500-RL-005) was carried out after a 24 h resting period following plating. Following cytokine stimulation, the cells were either collected for RNA isolation, Western blotting or were differentiated. To differentiate the cells, the culture medium was exchanged to medium without growth factors but supplemented with 1% fetal calf serum (FCS, Sigma Aldrich). The cells were differentiated for 5 days followed by fixation or preparation for WB or RNA isolation. Importantly, all experiments were performed on NSC after their second passage to ensure culture purity. Moreover, most experiments were performed or validated using NSCs cultures from wild type rats to ensure that the transgenic construct is not altering the cells in covert ways.

### RNAi

Plate-attached NSCs were transfected with 10μM of siRNA pools (ON-TARGETplus siRNA-SMARTpool, Dharmacon) using the RNAiMax system (Invitrogen, #13778100). The siRNA pools used were: rat ATF4: L-099212-02-0005; rat Ddit3: L-088282-02-0005; rat eIF2ak4: L-092044-02-0005; rat eIF2ak2: L-080001-02-0005; rat eIF2ak3: L-092012-02-0005 and the non-target control ON-TARGETplus non-targeting pool: D-001810-10-05. The medium was changed 24 hrs post-transfection and the cells were rested for one additional day before stimulation with cytokines, see previous section. For the expression of short hairpin RNA (shRNA) we used lentiviral plasmids pLKO.1-ATF4 for ATF4 (TRCN0000301721) with the target sequence: CGGACAAAGATACCTTCGAGT (#SHCLND, Merck) and for non-targeting controls a pLKO.1-plasmid encoding a scrambled sequence (shNTC). For the lentiviral production, HEK-293T cells were transfected using Lipofectamine 3000 (Invitrogen) with 500ng of the psPAX2, pMD2.G and the shATF4 containing plasmid.

Following 24 hrs, the media was changed to stem cell media without antibiotics. The virus-containing media were collected after additional 72 hrs, spinned down at 500Xg for 5 minutes and then filtered through a 0.45um filter (Sarstedt). Virus batches were always tested for proper ATF4 knockdown in the B16.F10 cell line. For transduction, the NSCs cultured into PDL-coated plates without antibiotics were transduced with 10% of either the shATF4 or shNTC lentivirus preparations for 24 hours. Their media was then changed, the cells were rested for additional 24 hours before selection with puromycin (1.5ug/ml) for four days and then used for experiments.

### Real-time qPCR

*Table 1* comprises all primer sequences and PCR programs used in this work. The samples and standards were run in duplicates by using a CFX384 touch Real-Time PCR detection system (BioRad) using SybrGreen-based (iQ™ SYBR® Green Supermix, BioRad) detection. Data analysis (amplification efficiency, melting curves and Ct values) were acquired by using the CFX Maestro software (BioRad). ΔCt values and expression values (2^^-ΔCt^) were calculated manually by using βactin as a house-keeping gene. The results were validated using 18S.

**Table 1.**
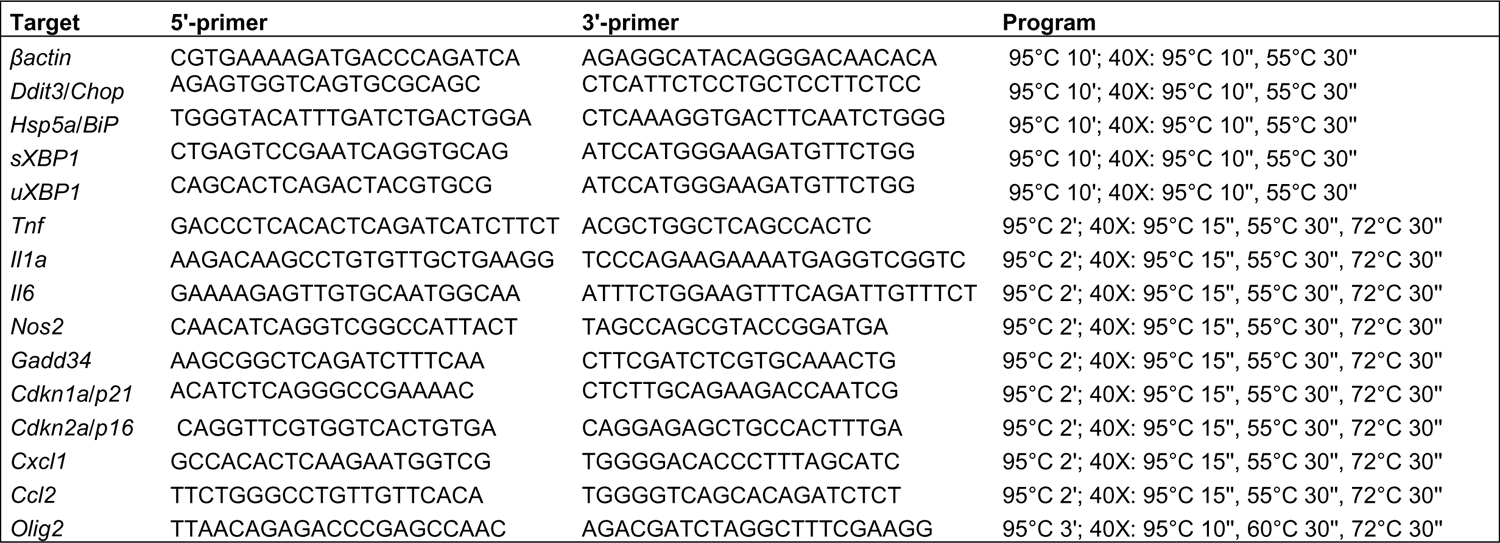
Primer sequences for real-time RT-PCR.

### CUT&RUN

200,000 primary NSCs either stimulated with IL1α (10ng/ml) or left unstimulated were used for CUT&RUN assay using the CUT&RUN kit from Cell Signaling Technologies® #86652 according to the manufacturer’s protocol. The following antibodies, all from Cell Signaling Technologies®, were used for targeted digestion of the positive control, isotype control and ATF4, respectively: Tri-methyl-Histone H3 (lys4) (C42D8) rabbit mAb (2μl/sample) Rabbit (DA1E) mAb IgG XP® isotype control (CUT&RUN) #66362 (5μl/sample) and rabbit anti-ATF4 antibody #11815S (2μl per sample). For sample normalization, 10μl of spike-in DNA (#36598, 1ng/μl) was added to each reaction. Following DNA isolation and purification using DNA purification and Spin columns from Cell signaling (ChiP, CUT&RUN) #14209, the sequencing libraries were generated using a NEBnext® Ultra™ II DNA library prep kit and NEBNext Multiplex Oligos (NEB #E7335) for Illumina (New England Biolabs). For size selection (25-150bp) and purification of the adaptor-ligated DNA and amplified library, SPRIselect beads were used (Beckman Coulter, Inc. #B23317). The sequencing libraries quantification and determination of size distribution was performed using an Agilent Bioanalyzer high Sensitivity DNA chip (Agilent). Qubit 3.0 using the Qubit dsDNA High Sensitivity Assay kit (Thermo Fisher Scientific) followed by equimolar pooling of the indexed libraries. The pooled library was sequenced using Illumina NextSeq 550 with 2×75 bp paired-end reads. Quality of reads were analyzed with FastQC, and STAR alignment (-alignIntronMax 1) was used to generate bam files. Peak detection was performed using SEACR (PMID: 31300027). Signals were displayed in Integrated Genomics Viewer (IGV;[28]).

### Seahorse Real-Time Cell Metabolic Analysis

The Oxygen Consumption rates (OCR) and Extracellular Acidification Rate (ECAR) were measured by using a SeaHorse XF96. NSCs were seeded into a PDL-coated XF96 cell culture microplate at 22,000 cells/well in 200ul propagation medium without antibiotics (see NSC culture section). The cells were transduced for 24hrs with lentiviral plasmids encoding either an ATF4 (shATF4) or scrambled control (shNTC) sequence. After removal of the virus and further allowing for additional 24h of resting, transduced NSCs were selected using puromycin (1.5 ug/ml) for three days and then stimulated with IL-1α (1 ng/ml) under continued puromycin selection. For the Fatty Acid Oxidation (FAO) assay n=4 cultures were used, each from individual rats, while for the glucose assay the n=8. NSCs were cultured in assay media (supplemented with 5mM glucose and palmitate-BSA for the FAO assay and 25mM glucose for the glucose assay). One hour before either of the measurements the NSCs were kept in at 37°C without CO_2_. The OCR and ECAR were recorded at basal levels and following application of oligomycin (1μM), carbonyl-cyanide 4-trifluoromethoxy-phenylhydrazone (FCCP) (1.0–1.5μM). The running protocol was 3 min of mixing, 2 min of waiting and 3 min of measurement. All chemicals were purchased from Sigma-Aldrich if not otherwise stated. Data were normalized on the number of cells per well. Cell number quantification was done by staining nuclei with Hoescht 33342 (Molecular Probes) for 10 min and then imaging each well using BD pathway 855 (BD Biosciences,Franklin Lakes, U.S.) with 20×objective and montage 5×4. Cell number was counted with Cell profiler software.

### Intracisternal injections

Animals were kept sedated using inhalation with isoflurane. The rats were placed so that their head would tip forward and facilitate the injection into the cisterna magna. The injection was performed using a dental 27G needle (Terum0, DN-2721) bent at a 45-degree angle connected to polyethylene tubing and a Hamilton syringe. The injection area was disinfected using ethanol and the needle was introduced 7.5mm deep through the skin into the cisterna magna. A 10ul solution containing the Il-1α was injected slowly over a period of 10 seconds.

## RESULTS

### Traumatic brain injury induces cellular stress in rat adult neural stem cells

The rat offers an excellent disease model and has been characterized extensively for autoimmune inflammation and traumatic injuries [29–31]. Throughout our studies, we have used the Dark Agouti (DA) strain for exploring the effects of neuroinflammation on the biology of adult NSCs [32–34]. To facilitate the exploration of NSC in this model, we generated a *Sox2* reporter rat (RedSox2) with DsRedExpress2 (DsRed) controlled by the *Sox2* promoter (+ 6kb - TSS) and the SRR2 enhancer on the DA/HanRj background (Figure 1A).

The DsRed expression recapitulated the endogenous expression of *Sox2* in the SVZ (Figure 1B), and SVZ-derived NSC neurosphere cultures showed varied levels of expression of DsRed recapitulating the well-known varied expression of *Sox2* expression among NSCs (Supplementary Figure 1A). FACSorting of RedSox2-derived NSCs into DsRed positive (DsRed^pos^) and negative (DsRed^neg^) populations showed that the DsRed^neg^ cells form much smaller neurospheres, (Supplementary Figure 1A). Also, the distribution of DsRed^pos^ vs DsRed^neg^ cells in neurosphere cultures changed from almost equal levels 55.3 ±6.8% vs 44.7±6.8%, respectively, after the first culture passage, to a more DsRed^pos^ abundant culture (85.3±4.3%) versus 14.8 ±4.3% DsRed^neg^ after the second passage (Supplementary Figure 1B). This suggests that the DsRed^pos^ cells accumulate after the first culture passage, while DsRed^neg^ cells, either have a slower proliferation cycle or are starting to express DsRed contributing to the DsRed^pos^ population. NCSs have the capability of differentiating into the main three cell types in the CNS, neurons, astrocytes and oligodendrocytes. When investigating their differentiation, we did not detect any differences in astrocyte percentages between the DsRed populations, 86,7±6.1% in DsRed^pos^ and 90±5.8% in DsRed^neg^ Supplementary Figure 1C-1D). Neurogenesis was equal in DsRed^pos^ 0.3±0.2% vs DsRed^neg^ 0.3±0.2% cultures (Supplementary Figure 1E), but oligodendrogenesis differed, where DsRed^neg^ cells (11.4±3.4%) generated a higher oligodendrocyte percentage than their DsRed^pos^ counterparts (1.2±0.6%), which was similar to NSCs from wild type rats, 1.7±0.8% (Supplementary Figure 1G-H). The high gliogenicity and decrease in cell numbers after the first passage could imply a more fated and less stem-cell like nature of the DsRed^neg^ population. Altogether, these results point to the accuracy of the reporter identifying the neural stem cell population within the DsRed^pos^ cell fraction.

To examine the role of TBI-induced neuroinflammation on the SVZ NSCs, we induced severe TBI in RedSox2 rats by employing the controlled cortical impact (CCI) injury model [35]. The mechanical injury affects the cortex and the lateral ventricle adjacent to the injury site, but even sites distal to the injury will be affected due to the action of the “glymphatic system” [36]. The effect of TBI on SVZ NSCs was then assessed by transcriptomics using *ex vivo* sorted NSCs (DsRed^pos^) from the SVZ at days 7 and 28 post-injury, and from the injury site at day 28 (Figure 1C). Brains were also collected for immunocytochemistry analysis on day 7 and day 28 post-injury. CSF was collected for cytokine/chemokine quantification and used as a benchmark for overall inflammatory status of the TBI model (Figure 1C). Gene set enrichment analysis (GSEA) of the transcriptomic data from NSCs day 7 post-TBI, revealed that the biological functions most enriched in TBI-derived cells were related to cell cycle control and inflammatory response (Figure 1D). At day 28 post-TBI, (Figure 1E) and in NSCs isolated from the injury site (Figure 1F), we observed that more functions overall were enriched in TBI-isolated NSCs compared to NSCs from healthy rats and compared to the earlier time-point, (Table 2). As at the previous time-point, the strongest enriched biological functions pertained to inflammatory response and cell cycle regulation, but also stress/apoptosis-related functions appeared, such as UV response, protein secretion and the p53 pathway (Table 2). In addition, an induction of the unfolded protein response (UPR) was detected in NSCs from the injury site 28 days post-injury versus control NSCs (Figure 1G).

**Table 2.**
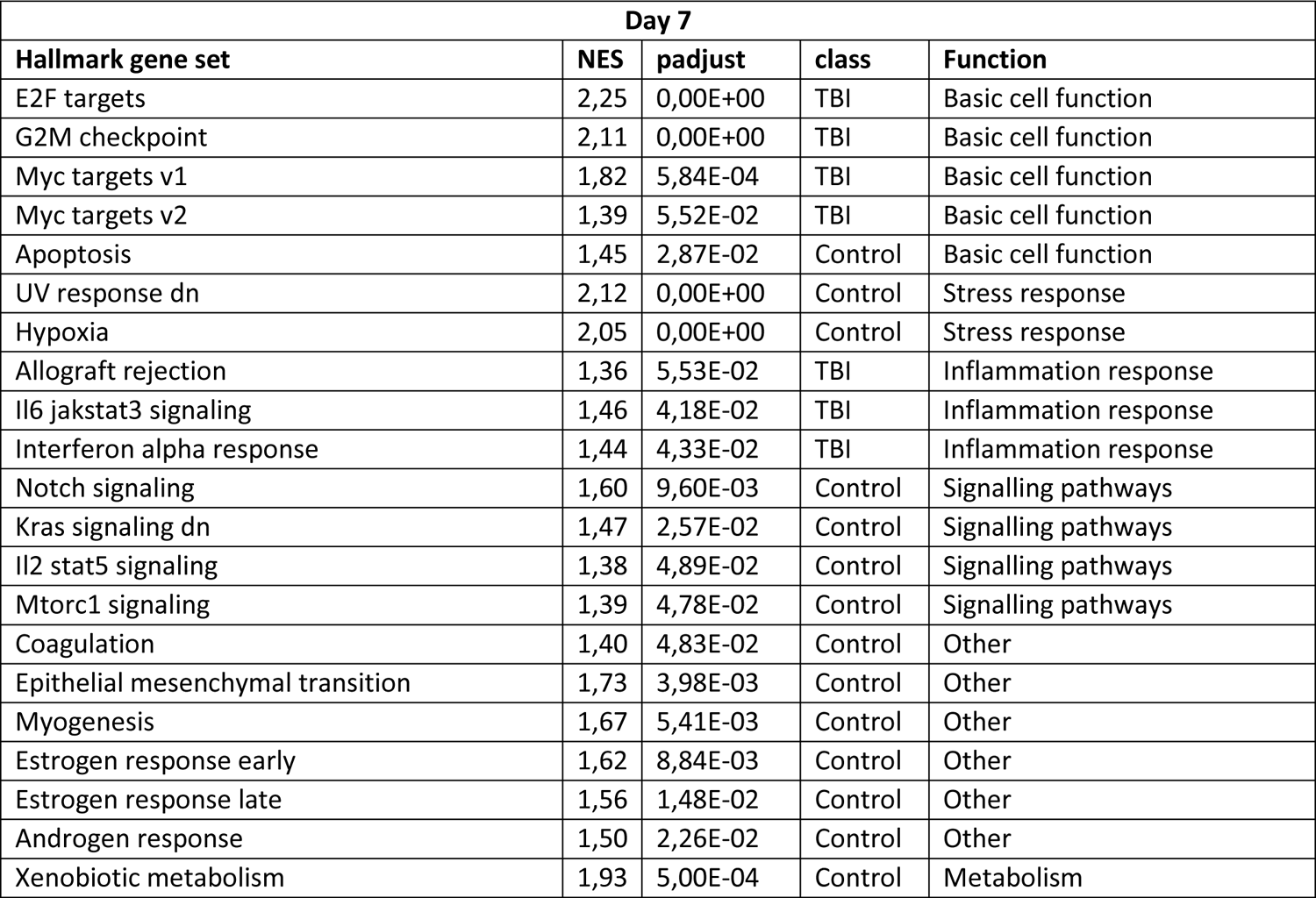
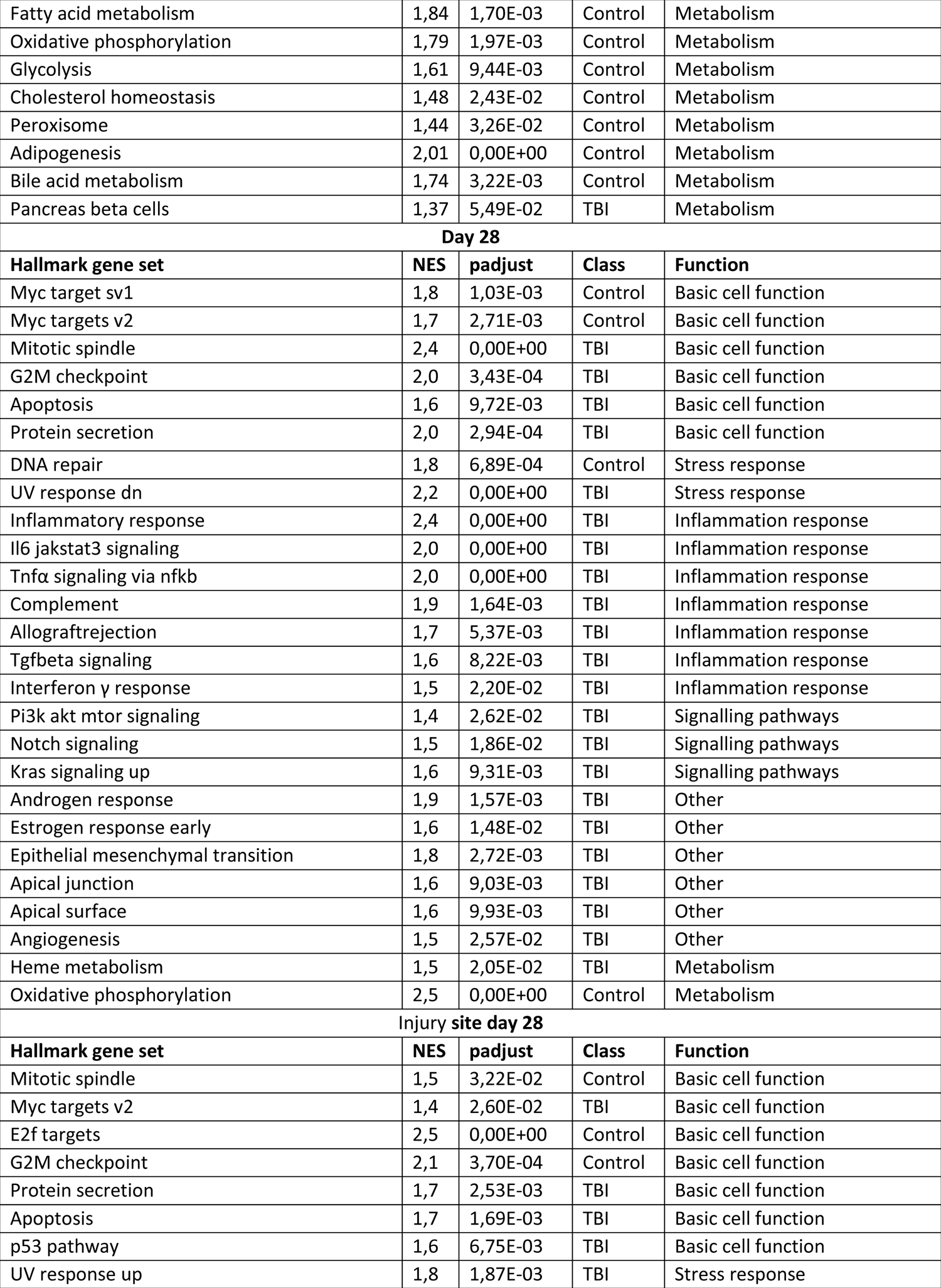
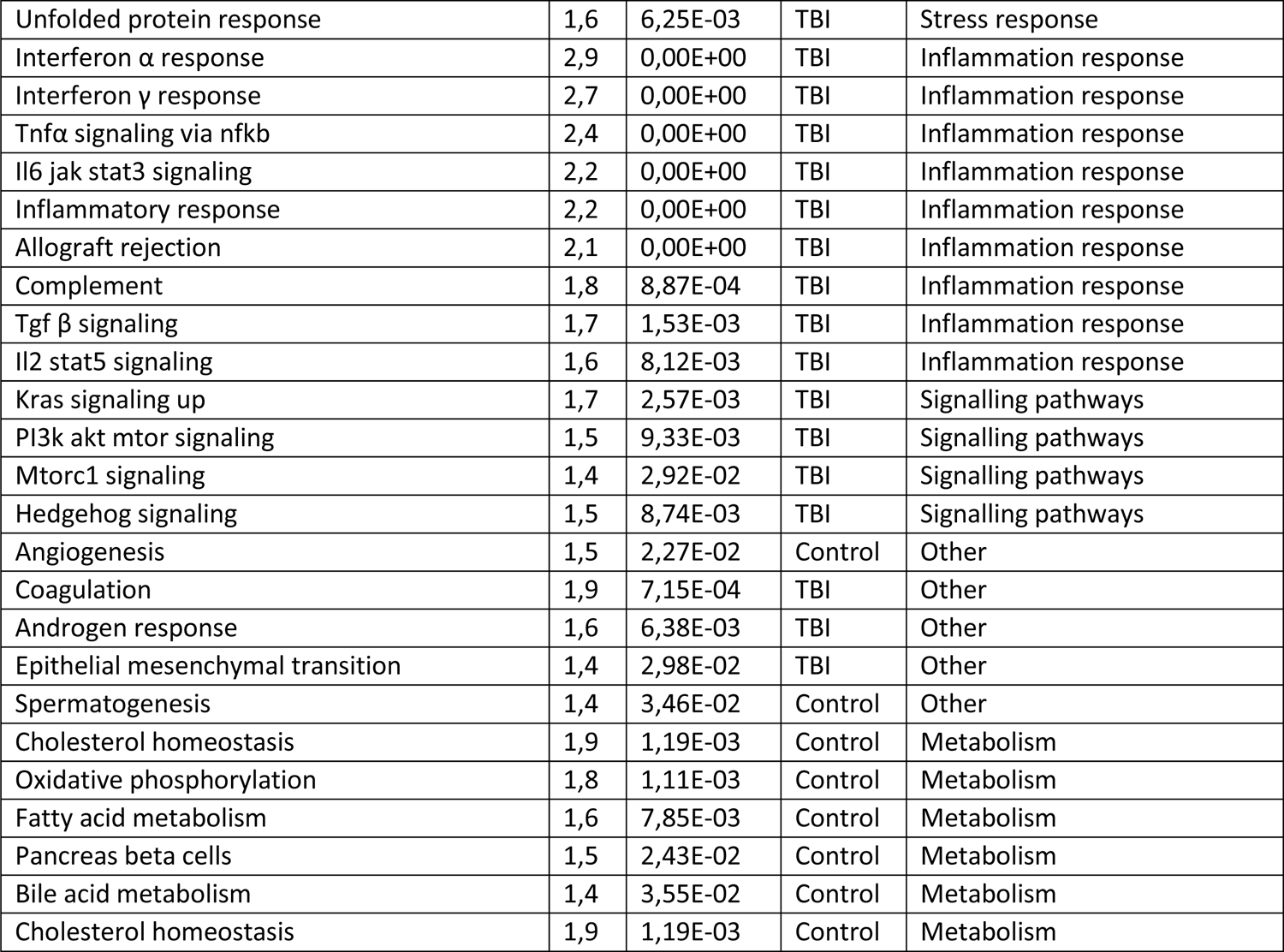
Hallmark gene sets significantly enriched in NSC from TBI vs Control rats and used to make the bubble plots in Figure 1. The column “class” shows in which experimental group the respective gene set was enriched.

| Day 7 |  |  |  |  |
| --- | --- | --- | --- | --- |
| Hallmark gene set | NES | padjust | class | Function |
| E2F targets | 2,25 | 0,00E+00 | TBI | Basic cell function |
| G2M checkpoint | 2,11 | 0,00E+00 | TBI | Basic cell function |
| Myc targets v1 | 1,82 | 5,84E-04 | TBI | Basic cell function |
| Myc targets v2 | 1,39 | 5,52E-02 | TBI | Basic cell function |
| Apoptosis | 1,45 | 2,87E-02 | Control | Basic cell function |
| UV response dn | 2,12 | 0,00E+00 | Control | Stress response |
| Hypoxia | 2,05 | 0,00E+00 | Control | Stress response |
| Allograft rejection | 1,36 | 5,53E-02 | TBI | Inflammation response |
| Il6 jakstat3 signaling | 1,46 | 4,18E-02 | TBI | Inflammation response |
| Interferon alpha response | 1,44 | 4,33E-02 | TBI | Inflammation response |
| Notch signaling | 1,60 | 9,60E-03 | Control | Signalling pathways |
| Kras signaling dn | 1,47 | 2,57E-02 | Control | Signalling pathways |
| Il2 stat5 signaling | 1,38 | 4,89E-02 | Control | Signalling pathways |
| Mtorc1 signaling | 1,39 | 4,78E-02 | Control | Signalling pathways |
| Coagulation | 1,40 | 4,83E-02 | Control | Other |
| Epithelial mesenchymal transition | 1,73 | 3,98E-03 | Control | Other |
| Myogenesis | 1,67 | 5,41E-03 | Control | Other |
| Estrogen response early | 1,62 | 8,84E-03 | Control | Other |
| Estrogen response late | 1,56 | 1,48E-02 | Control | Other |
| Androgen response | 1,50 | 2,26E-02 | Control | Other |
| Xenobiotic metabolism | 1,93 | 5,00E-04 | Control | Metabolism |
| Fatty acid metabolism | 1,84 | 1,70E-03 | Control | Metabolism |
| Oxidative phosphorylation | 1,79 | 1,97E-03 | Control | Metabolism |
| Glycolysis | 1,61 | 9,44E-03 | Control | Metabolism |
| Cholesterol homeostasis | 1,48 | 2,43E-02 | Control | Metabolism |
| Peroxisome | 1,44 | 3,26E-02 | Control | Metabolism |
| Adipogenesis | 2,01 | 0,00E+00 | Control | Metabolism |
| Bile acid metabolism | 1,74 | 3,22E-03 | Control | Metabolism |
| Pancreas beta cells | 1,37 | 5,49E-02 | TBI | Metabolism |
| **Day 28** |  |  |  |  |
| **Hallmark gene set** | **NES** | **padjust** | **Class** | **Function** |
| Myc target sv1 | 1,8 | 1,03E-03 | Control | Basic cell function |
| Myc targets v2 | 1,7 | 2,71E-03 | Control | Basic cell function |
| Mitotic spindle | 2,4 | 0,00E+00 | TBI | Basic cell function |
| G2M checkpoint | 2,0 | 3,43E-04 | TBI | Basic cell function |
| Apoptosis | 1,6 | 9,72E-03 | TBI | Basic cell function |
| Protein secretion | 2,0 | 2,94E-04 | TBI | Basic cell function |
| DNA repair | 1,8 | 6,89E-04 | Control | Stress response |
| UV response dn | 2,2 | 0,00E+00 | TBI | Stress response |
| Inflammatory response | 2,4 | 0,00E+00 | TBI | Inflammation response |
| Il6 jakstat3 signaling | 2,0 | 0,00E+00 | TBI | Inflammation response |
| Tnfα signaling via nfkb | 2,0 | 0,00E+00 | TBI | Inflammation response |
| Complement | 1,9 | 1,64E-03 | TBI | Inflammation response |
| Allograftrejection | 1,7 | 5,37E-03 | TBI | Inflammation response |
| Tgfbeta signaling | 1,6 | 8,22E-03 | TBI | Inflammation response |
| Interferon γ response | 1,5 | 2,20E-02 | TBI | Inflammation response |
| Pi3k akt mtor signaling | 1,4 | 2,62E-02 | TBI | Signalling pathways |
| Notch signaling | 1,5 | 1,86E-02 | TBI | Signalling pathways |
| Kras signaling up | 1,6 | 9,31E-03 | TBI | Signalling pathways |
| Androgen response | 1,9 | 1,57E-03 | TBI | Other |
| Estrogen response early | 1,6 | 1,48E-02 | TBI | Other |
| Epithelial mesenchymal transition | 1,8 | 2,72E-03 | TBI | Other |
| Apical junction | 1,6 | 9,03E-03 | TBI | Other |
| Apical surface | 1,6 | 9,93E-03 | TBI | Other |
| Angiogenesis | 1,5 | 2,57E-02 | TBI | Other |
| Heme metabolism | 1,5 | 2,05E-02 | TBI | Metabolism |
| Oxidative phosphorylation | 2,5 | 0,00E+00 | Control | Metabolism |
| **Injury site day 28** |  |  |  |  |
| **Hallmark gene set** | **NES** | **padjust** | **Class** | **Function** |
| Mitotic spindle | 1,5 | 3,22E-02 | Control | Basic cell function |
| Myc targets v2 | 1,4 | 2,60E-02 | TBI | Basic cell function |
| E2f targets | 2,5 | 0,00E+00 | Control | Basic cell function |
| G2M checkpoint | 2,1 | 3,70E-04 | Control | Basic cell function |
| Protein secretion | 1,7 | 2,53E-03 | TBI | Basic cell function |
| Apoptosis | 1,7 | 1,69E-03 | TBI | Basic cell function |
| p53 pathway | 1,6 | 6,75E-03 | TBI | Basic cell function |
| UV response up | 1,8 | 1,87E-03 | TBI | Stress response |
| Unfolded protein response | 1,6 | 6,25E-03 | TBI | Stress response |
| Interferon $\alpha$ response | 2,9 | 0,00E+00 | TBI | Inflammation response |
| Interferon $\gamma$ response | 2,7 | 0,00E+00 | TBI | Inflammation response |
| Tnfα signaling via nfkb | 2,4 | 0,00E+00 | TBI | Inflammation response |
| Il6 jak stat3 signaling | 2,2 | 0,00E+00 | TBI | Inflammation response |
| Inflammatory response | 2,2 | 0,00E+00 | TBI | Inflammation response |
| Allograft rejection | 2,1 | 0,00E+00 | TBI | Inflammation response |
| Complement | 1,8 | 8,87E-04 | TBI | Inflammation response |
| Tgf $\beta$ signaling | 1,7 | 1,53E-03 | TBI | Inflammation response |
| Il2 stat5 signaling | 1,6 | 8,12E-03 | TBI | Inflammation response |
| Kras signaling up | 1,7 | 2,57E-03 | TBI | Signalling pathways |
| PI3k akt mtor signaling | 1,5 | 9,33E-03 | TBI | Signalling pathways |
| Mtorc1 signaling | 1,4 | 2,92E-02 | TBI | Signalling pathways |
| Hedgehog signaling | 1,5 | 8,74E-03 | TBI | Signalling pathways |
| Angiogenesis | 1,5 | 2,27E-02 | Control | Other |
| Coagulation | 1,9 | 7,15E-04 | TBI | Other |
| Androgen response | 1,6 | 6,38E-03 | TBI | Other |
| Epithelial mesenchymal transition | 1,4 | 2,98E-02 | TBI | Other |
| Spermatogenesis | 1,4 | 3,46E-02 | Control | Other |
| Cholesterol homeostasis | 1,9 | 1,19E-03 | Control | Metabolism |
| Oxidative phosphorylation | 1,8 | 1,11E-03 | Control | Metabolism |
| Fatty acid metabolism | 1,6 | 7,85E-03 | Control | Metabolism |
| Pancreas beta cells | 1,5 | 2,43E-02 | Control | Metabolism |
| Bile acid metabolism | 1,4 | 3,55E-02 | Control | Metabolism |
| Cholesterol homeostasis | 1,9 | 1,19E-03 | Control | Metabolism |

Overall, TBI had a dramatic effect on the SVZ NSCs by activating pathways related to inflammation and cell cycle progression and suggesting that factors from the injury site could affect the SVZ NSC niche. Moreover, the UPR was triggered in NSC from the injury site.

### Traumatic brain injury activates the integrated stress response (ISR) in rat neural stem cells via IL-1α and ATF4

Next, we wanted to elucidate what activates the UPR, since it can be achieved by either the ER-stress or ISR pathways. Although the UPR has previously been associated to TBI [9] it is still unclear how UPR is triggered and what the consequences are to the NSCs. Since TBI induced transcriptional changes in NSCs, not only at the injury site but also in the SVZ, we hypothesized that soluble factors from the injury site would influence the NSCs in the SVZ. To test this, we quantified TBI-induced factors in the cerebral spinal fluid (CSF) using multiplex ELISA. We discovered a strong increase of IL-1α in the CSF at day 7 post-injury, (Figure 2A), and a strong enrichment of IL-1α-regulated genes in NSCs isolated from the injury site of TBI rats (Figure 2B). To determine if IL-1α could induce the ER-stress in NSCs, we exposed the cells to various concentrations of IL-1α followed by quantification of ER-stress/UPR signature transcripts and the spliced isoform of the canonical ER stress transcription factor *Xbp1*. Since IL-1α did not significantly induce splicing of *Xbp1* (Figure 2C), we reasoned that the activation of UPR signature genes was driven mainly by the ISR and not canonical ER stress. Indeed, the master regulator of ISR, ATF4, was significantly induced by IL-1α (Figure 2D and 2E). These results suggest that the induction of UPR genes was caused by the ISR following exposure to IL-1α. Importantly, canonical target genes of ATF4 (*Hspa5/BiP Gadd34* and *Ddit3/Chop*), were induced in NSCs by concentrations of IL-1α equal to those detected in the CSF of TBI rats (Figure 2A and Supplementary Figure 2A-C). Moreover, the activation of these target genes was abrogated by ATF4 knockdown (Figure 2F, 2G, and 2H). To verify that ATF4 was expressed in NSCs *in vivo* after TBI, we stained for ATF4 in coronal sections of TBI brains (Figure 2N-P). ATF4 expression was detected in TBI brains, close to the ventricle (Figure 2N-P) and in the injury area (not shown). Next, we wanted to elucidate how IL-1α induced activation of ATF4 translation. Since the classical signaling pathway leading to ATF4 translation is mediated via the phosphorylation of eIF2α [37] we tested the effect of IL-1α on eIF2α phosphorylation. Indeed, NSCs exposed to IL-1α had an increased level of phosphorylated eIF2α and increased levels of ATF4 (Figure 2I). This could be blocked by pretreatment with a small molecule inhibitor of eIF2α (ISRIB) (Figure 2I-K). Next, we profiled the genome-wide occupancy of ATF4 after IL-1α stimulation using CUT&RUN-sequencing (Figure 2L, 2M). IL-1α induced a global increase of ATF4 binding to transcription start sites (TSS) and we confirmed that IL-1α induced binding of ATF4 to TSSs of several UPR or ER-associated protein degradation (ERAD)-associated genes, exemplified by *Derl1* and *Edem1*, respectively. The occupancy density graphs for these two genes are shown in (Figure 2L-M). In summary, IL-1α activates eIF2α in NSCs, which leads to translation of ATF4, which in turn initiates the expression of its canonical target genes *Hspa5/BiP, Gadd34* and *Ddit3/Chop*.

**Figure 2.**
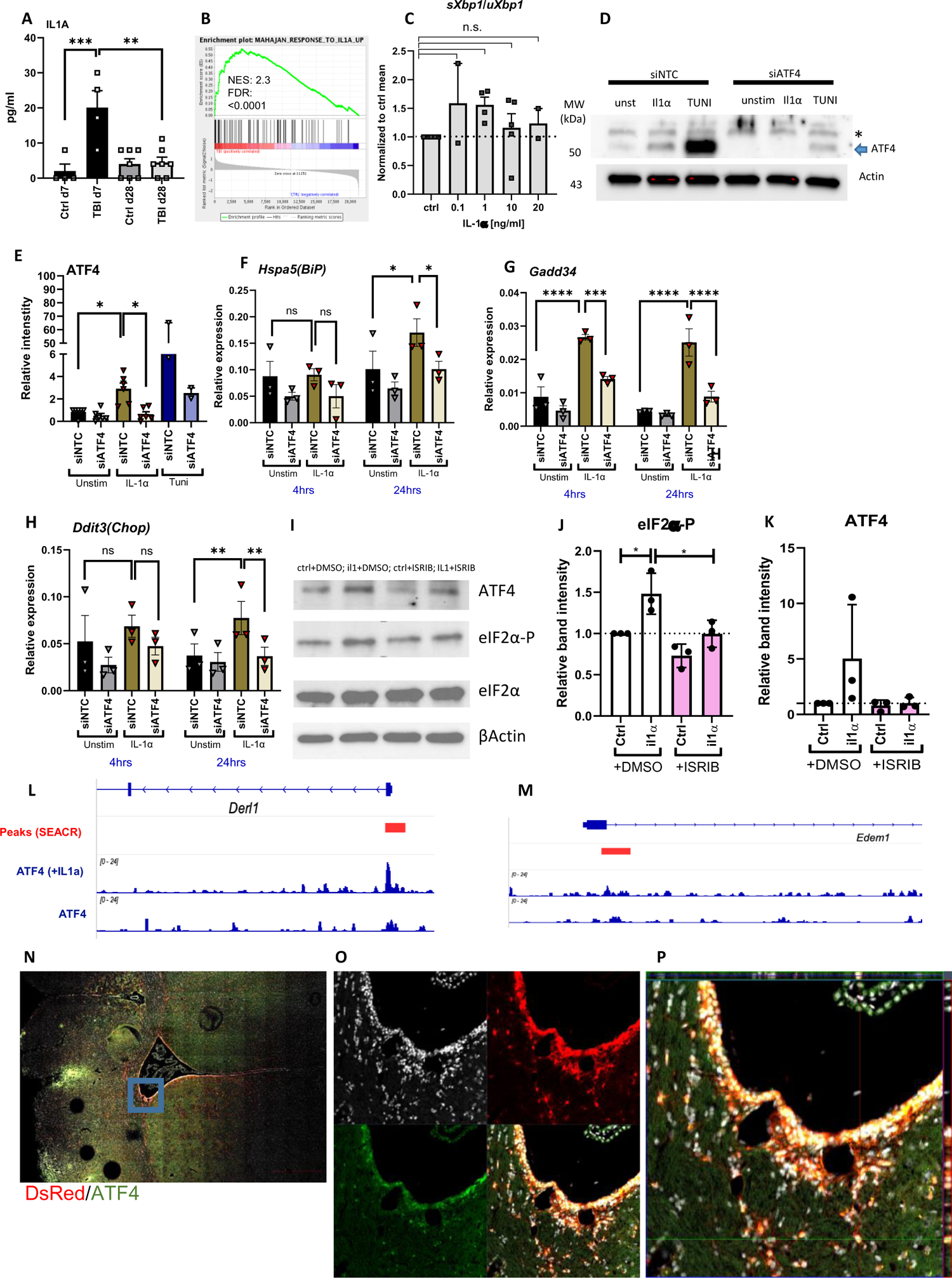
IL-1a induces ISR/ATF4 pathway in NSCs. (**A**). IL-1a ELISA in CSF collected from rats with TBI 7 and 28 days post-injury. (**B**). Enrichment plot showing enrichment of IL-1a response gene set in NSC isolated from TBI rats. NES=normalized enrichment score, FDR=false discovery rate. (**C**). Graph show levels of expression ratio between spliced XBP-1 and unspliced XBP-1 following stimulation with IL-1a cytokine for 24hrs. The data is shown as normalized to unstimulated control, n=2-4. **D**).Representative pictures of immunoblotting against ATF4 (blue arrow at ∼50kDa) in NSCs stimulated with IL-1a (24hrs) or Tunicamycin (Tuni, 5hrs) or kept unstimulated (unst) after KD with siRNA for ATF4 (siATF4) or non-targeting control (siNTC). *=non-specific band. (**E**) Quantification of band intensities from WB n=2-5. (F-H) Real-time RT-PCR quantification of RNA expression normalized to actin of Hspa5(Bip) (**F**) Gadd34 (**G**) and Ddit3(Chop) (**H**) after 4h or 24hrs of stimulation with IL1a. (**I**)Representative pictures of immunoblotting against ATF4, phosphorylated and total eIF2a and βactin. (**J-K**) Quantification of WB n=3 for eIf2a - P (**J**) and ATF4 (**K**) after a 5h stimulation with IL-1a and/or DMSO/ISRIB. Data was normalized to total eIF2a and β-actin (**J**) and to β-actin (**K**) and presented as fold expression to unstimulated control (**L**). ATF4 occupancy at Derl1(**L**) and Edem1(**M**) in NSCs rat stimulated with IL-1a using the CUT&RUN sequencing. (**N-P**). Coronal sections of RedSox rat brain with TBI immunelabeled forATF4 (green). Overview picture of lateral ventricle (N). Blue square magnified in O and P. Orthogonal projection of DsRed^pos^ and ATF4^pos^cell (P). In all graphs bars show mean ±SEM; dots show biological replicates or individual experiments. Statistics: One way ANOVA, Tukey’s or Holm-Šídák’s multiple comparison test ***p>0.001; **p>0.01.

### ATF4 is necessary for IL-1α-induced transcriptional changes related to metabolism and inflammatory pathways

To determine what effects of IL-1α were mediated though ISR/ATF4 activation in NSCs, we stimulated NSCs with IL-1α (4 or 24 hours) after transfection with ATF4-specific siRNA or non-targeting control (NTC) siRNA and then performed RNA-sequencing (Figure 3A). Two major clusters were identified after IL-1α stimulation by using Multiple Experiment Viewer (MeV) and WEB-based Gene Set AnaLysis Toolkit (WEBGestalt, ref) (Figure 3B). Cluster 1 (downregulated genes) was enriched for genes associated to metabolic pathways such as carbon metabolism and fatty acid metabolism (Figure 3B and 3C). In contrast, cluster 2 (upregulated genes) was enriched for genes involved in inflammatory signaling pathways (TNF, NF-κB, JAK-STAT) as well as p53 and MAPK signaling (Figure 3D). Ingenuity pathway analysis (IPA) identified an upregulation of inflammatory pathways following IL-1α stimulation and the abrogation of this upregulation following ATF4 knockdown (Figure 3E).

**Figure 3.**
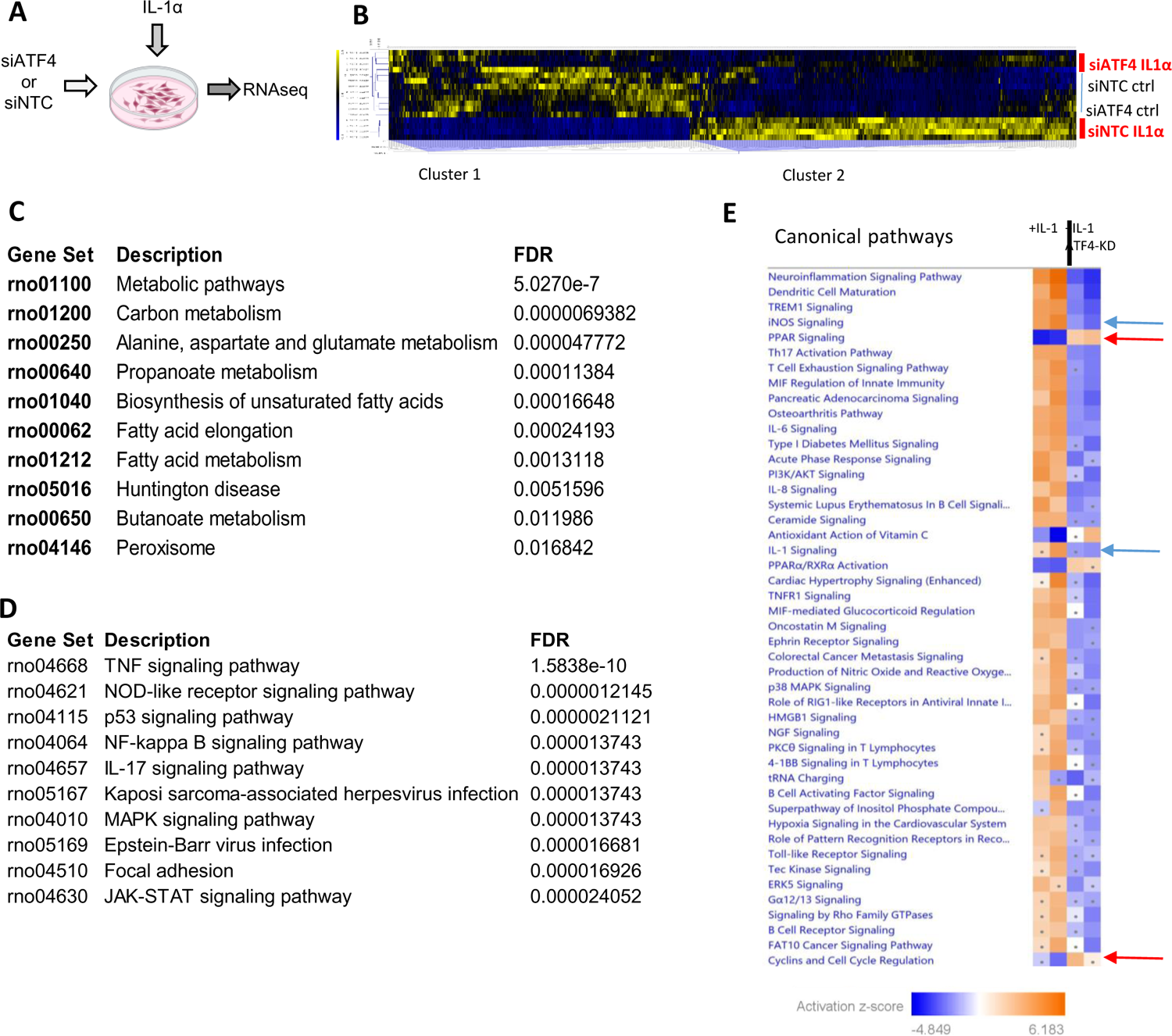
Functional annotation of RNAsequencing data obtained from NSCs with or without ATF4 knockdown and/or stimulated with IL1a for 4 or 24hrs. (**A**) Experimental set-up overview (created with BioRender.com), n=4 individual NSC cultures. (B) Heat map showing hierarchically clustered genes that are differentiatlly expressed between IL1a groups that are either knockdown for ATF4 or not. (**C-D**) Gene onthology analysis of cluster1 (**C**) and cluster 2(**D**) using the WEbGestalt tool. (**E**) Ingenuity pathway analysis showing significantly regulated canonical pathways. Blue= downregulated pathways, yellow=upregulated. Color intensity = streghth of z-score. Z-score>2 = significant.

### IL-1α via ATF4, shifts the metabolic activity of NSCs towards glycolysis

The alteration of metabolic gene expression in NSCs after IL-1α stimulation suggested that IL-1α affects the metabolic activity of NSCs. To verify this, we performed Seahorse analysis of NSCs stimulated with IL-1α and determined the oxygen consumption rates (OCR) and extracellular acidification rates (ECAR). To identify ATF4-dependent effects on NSC metabolism we transduced the cells with lentiviral plasmids encoding for NTC or ATF4-specific shRNA (Figure 4A). After transduction, the NSCs were selected with puromycin for four days and stimulated with IL-1α for the last 24 hours. OCR or ECAR were not affected when palmitate was used as the main energy source in the media (Supplementary figure S2D-F) suggesting that FAO (Fatty Acid Oxidation) was not affected. In contrast, when glucose was used as the energy source, we detected a significant decrease in basal OCR/ECAR ratio upon IL-1α stimulation, which was impaired by ATF4 knockdown (Figure 4B). When investigating the effects on OCR (Figure 4C) and ECAR (Figure 4D) individually, it was clear that the OCR/ECAR ratio change was attributed to an increase in ECAR. ECAR can be a proxy measurement for glycolysis, and the matching shift in lactate levels measured in the media (Figure 4D) strengthened that hypothesis. In all, we demonstrate that NSCs undergo a metabolic shift towards glycolysis after IL-1α stimulation, and this shift is mediated by ATF4.

**Figure 4.**
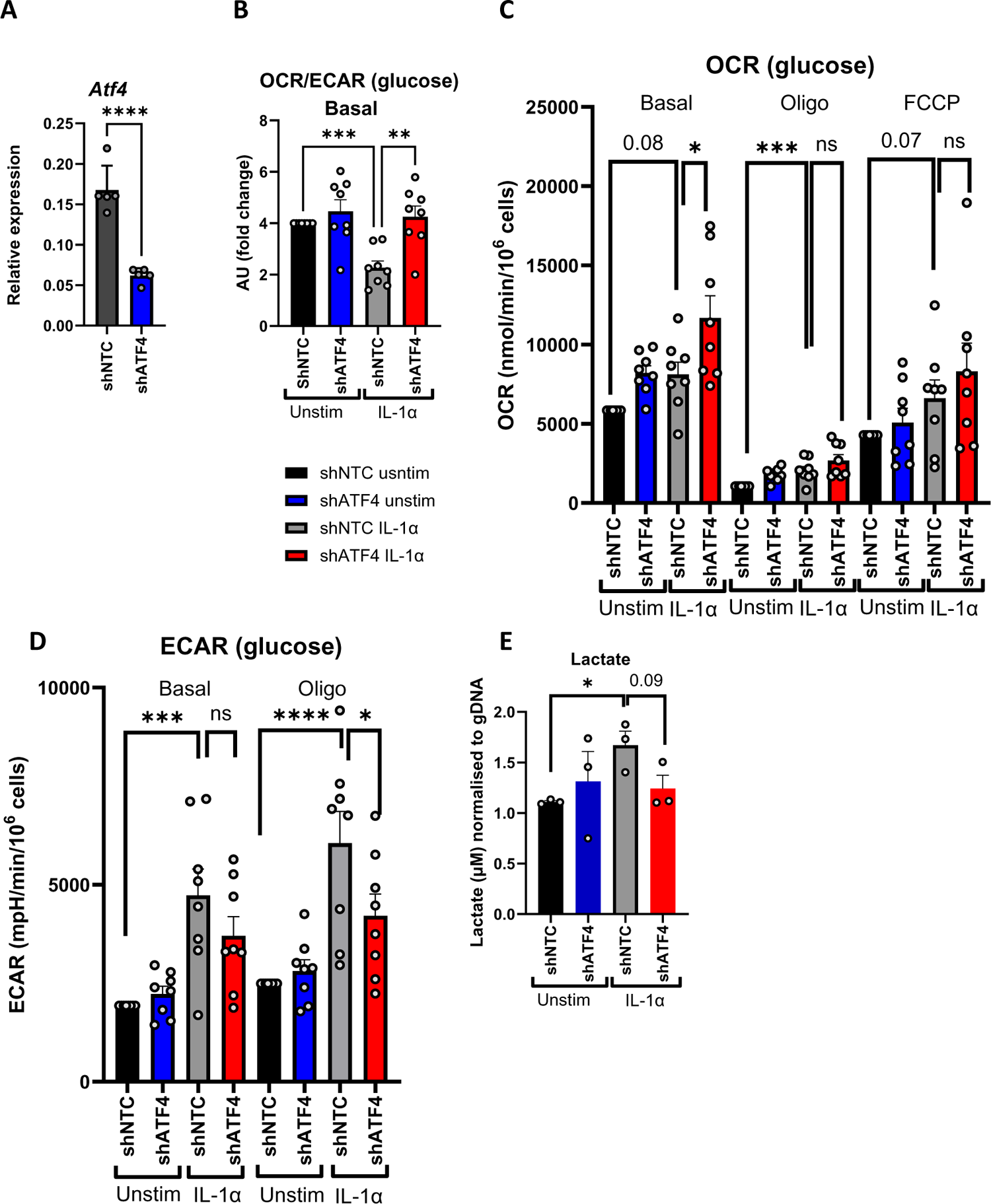
IL-1a induces an ATF4-mediated metabolic shift towards glycolysis. (**A**) RNA expression levels of ATF4 following lentiviral transduction with plasmids containing shRNA to ATF4 (shATF4) or scrambled control (shNTC). ATF4 expression values are normalized to βactin. NSCs were transduced with either constructs and following a four day selection with puromycin were stimulated with IL-1a or kept unstimulated (unstim) for 24 hours. Seahorse measurements were performed at the end of the 24h period. (**B**) The ratio between Oxygen Consumption Rate (OCR) and Extracellular Acidification Rate (ECAR) area under curve (AU) shown as fold change of unstimulated shNTC control. (**C**) OCR measured in nmol/min and normalized to number of NSCs. Values shown are normalized to unstimulated shNTC control. (**D**) ECAR measured in mpH/min and normalized to number of NSCs. Values shown are normalized to unstimulated shNTC. (**E**) Lactate levels (μM) measured in NSC supernatants exposed to the same conditions as in (A) normalized to gDNA as a proxy for total cell numbers, n=3. In all graphs bars show mean ±SEM; dots show biological replicates or individual experiments. Statistics: One way ANOVA, Tukey’s or Holm-Šídák’s multiple comparison test ***p>0.001; **p>0.01, *>0.05. Oligo = olygomycin 1μM; FCCP 1.0–1.5μM.

### IL-1α induces cell cycle arrest via ATF4

Since metabolic features are closely related to cellular functions such as cell proliferation, we measured the percentage of proliferating NSCs by using ki67 immunolabeling (Figure 5 A-D). Following IL-1α exposure, the percentage of ki67+ NSCs significantly decreased, and the effect could be reverted by ATF4-knockdown (Figure 5E). Moreover, CDKN1A/p21^Cip1^, a cyclin dependent kinase inhibitor, involved in maintaining the NSC pool by keeping the NSC in a quiescent state, [38] was upregulated by IL-1α both on transcriptional (Figure 5F) and at the protein level (Figure 5H, 5I) and its upregulation was hampered by ATF4-knockdown.

**Figure 5.**
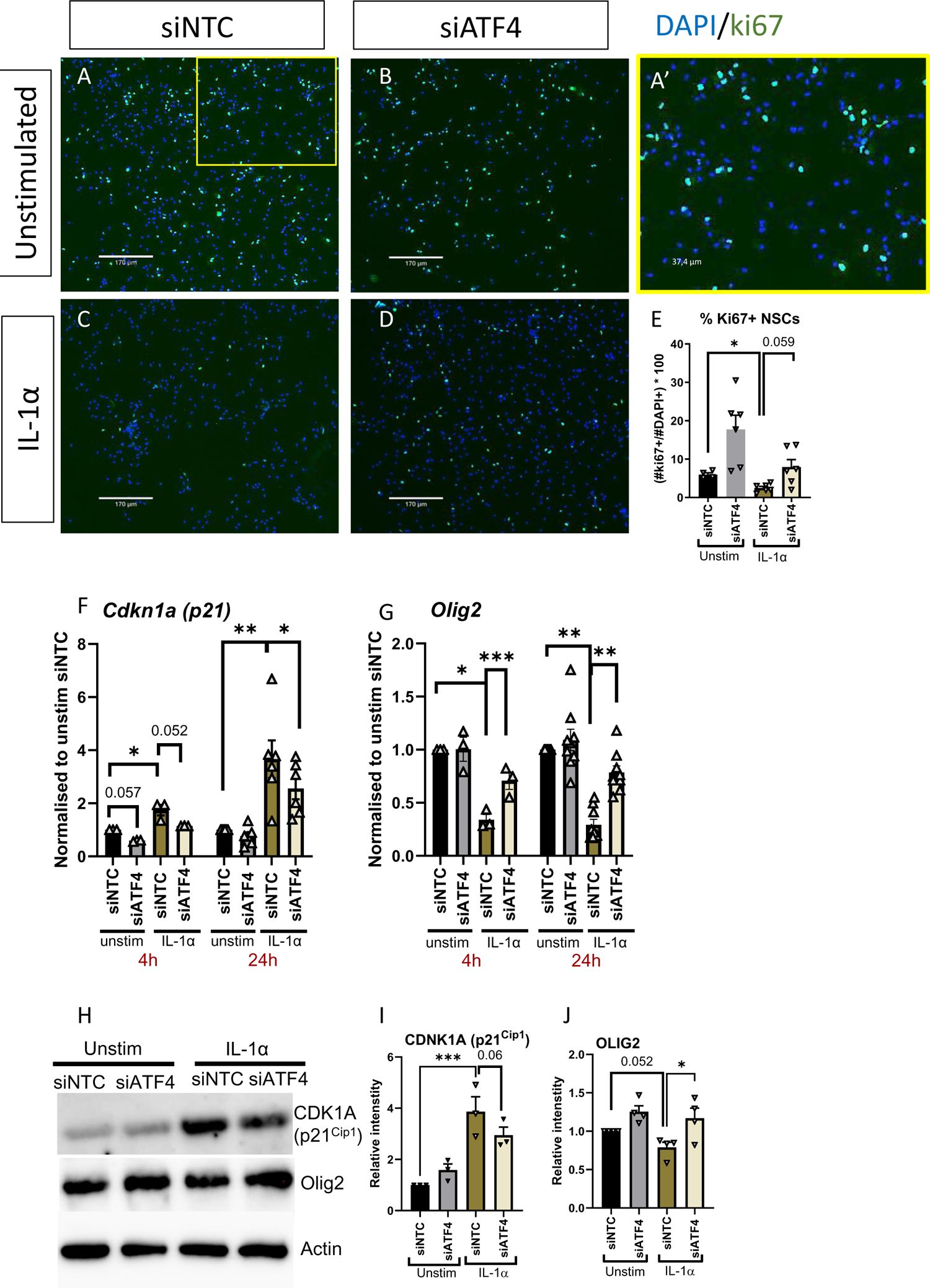
IL1a induces cell cycle arrest via ATF4. (**A-D**) Representative pictures of undifferentiated NSCs immunelabeled for Ki67 (green) or nuclear staining DAPI (blue). The NSCs had either been transfected with siRNA for NTC control (**A**), ATF4 (**B**) and kept unstimualted (**C**) or stimulated with IL-1a for 24hrs (**D**). (**A’**) The yellow-marked inlet in (A) is shown in a bigger magnification in A’. (**E**) Quantification of the percentage of Ki67+ NSC of total cell number given by DAPI+ nuclei. (**F-G**) Quantification of gene expression using real-time RT-PCR for Cdk1a(p21) (**F**) and Olig2(**G**). Raw values were normalised to βactin and plotted as fold change to unstimulated control (siNTC usntim). (**H**) Representative pictures of immunoblotting depicting bands specific for CDK1A(p21), Olig2 and βactin (**I,J**). Three WB images from different experiments were used to quantify the band intensity of the respictive bands CDK1A(p21) (**I**), OLIG2(**J**). The band intensities were normalised to βactin band intensities and were ploted as a fold change to unstimulated control (siNTC unstim). In all graphs bars show mean ±SEM; dots show biological replicates or individual experiments. Statistics: One way ANOVA, Tukey’s or Holm-Šídák’s multiple comparison test ***p>0.001; **p>0.01, *>0.05.

Interestingly, there was a reverse effect by IL-1α on OLIG2, a regulator of self-renewal in NSC. OLIG2 was strongly downregulated by IL-1α, both transcriptionally (figure 5G) and translationally (Figure 5H, 5J). Overall, IL-1α induces cell cycle arrest in NSC via CDK1A/p21^Cip1^ upregulation and OLIG2 downregulation which is mediated via ATF4.

### IL-1α /ATF4 axis induces senescence in NSCs

The RNA-sequencing data revealed that IL-1α lead to an increased pro-inflammatory gene expression signature in NSCs. RT-qPCR analysis of pivotal inflammatory genes, *Nos2*, *Tnf, and Il1a* confirmed that upon stimulation with IL-1α the expression of these genes was significantly upregulated in NSCs (Figure 6A, 6B, and Supplementary Figures 2I-L) and completely abrogated by ATF4 knockdown (Figure 6A, 6B). The secretion of TNF protein from IL-1α-stimulated NSCs followed the same pattern as the gene expression (Figure 6C). The expression of other inflammatory factors, such as IL-6 was not significantly regulated (not shown) and neither was that of the anti-inflammatory cytokine IL-10 (not shown). To test the effect of IL-1α *in vivo*, we performed intracisternal injection of IL-1α in RedSox2 rats, followed by NSC isolation and quantification of gene expression using RT-qPCR (Figure 6D-6I). Of the genes investigated, we could see a significant upregulation of *Tnf* (Figure 6H) and a tendency of increase in *Gadd34* (Figure 6I) following a single pulse with IL-1α and investigation after a 24h incubation period. On the other hand, in DsRed^neg^ cells, IL-1α stimulation had a strong reversed tendency of downregulation for *Tnf* (data not shown) and no effect on *Gadd34* (data not shown).

**Figure 6.**
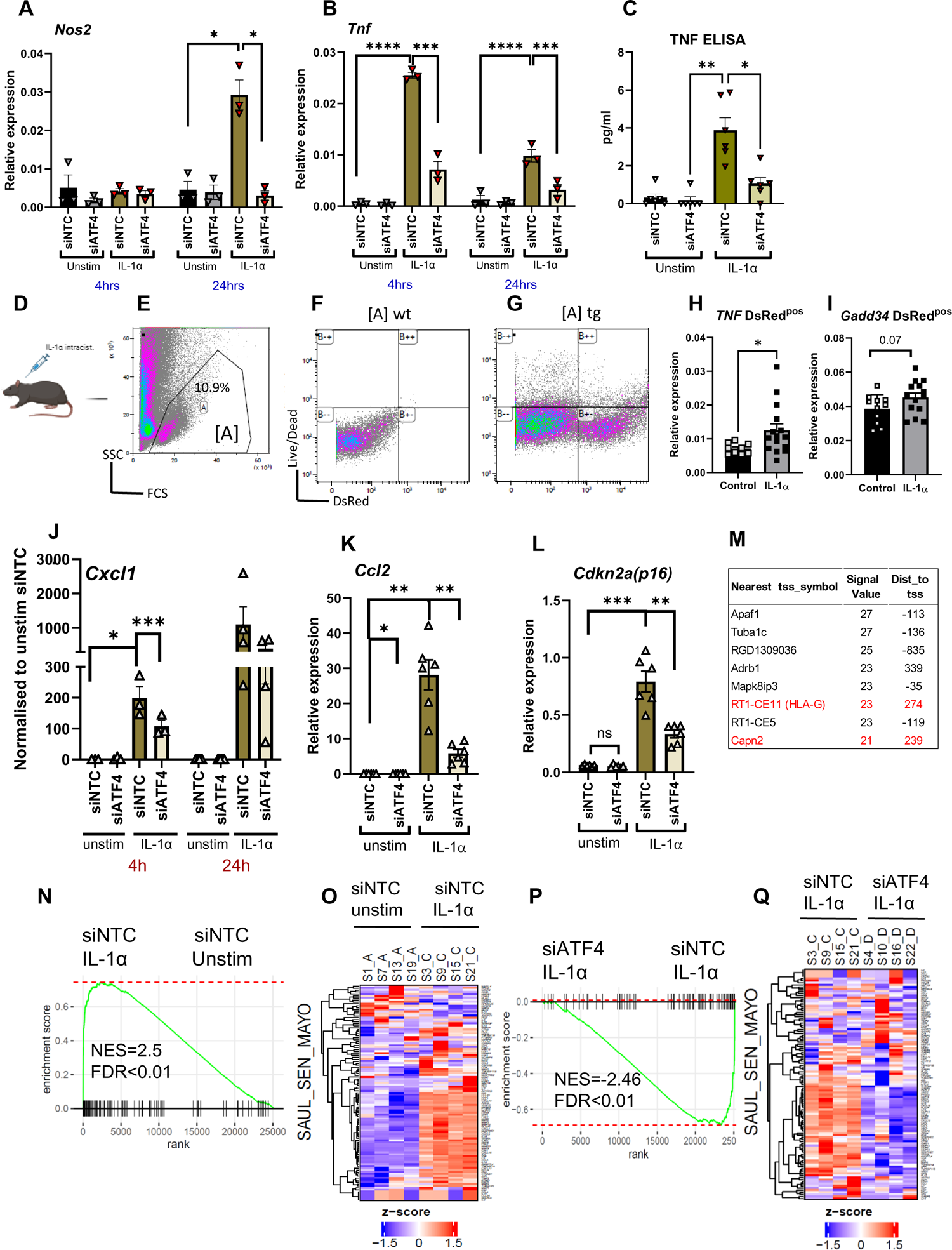
ATF4 is necessary for IL-1a-induced senescence in NSCs. (**A-C**) Quantification of gene expression using real-time RT-PCR for Nos2 (**A**) and Tnf (**B**). Raw values were normalised to βactin. The NSCs had been transfected with siRNA to control (siNTC) or ATF4 (siATF4) and/or stimulated with IL-1a for 4 or 24 hrs. (**C**)TNF measured in NSC culture supernatatnt using ELISA. (**D**) Schematic drawing of the intracisternal injection of IL-1a in RedSox rats. (**E-G**) Representative FACSoritng density plots showing the FSC/SSC population sorted (E) the widltype control, wt (F) and the RedSox, Red^neg^ and Red^pos^ populations (G). (**H-I**) Gene expression assessed with real-time RT-PCR in Red^pos^ of Tnf (**H**) and Gadd34 (**I**).(J-K) Gene expression assessed with real-time RT-PCR in NSC after treatment similar to A-C; Cxcl1 (**J**), Ccl2(**K**) and Cdkn2a(p16) (**L**). Values in A-C and J-L were nomalised to βactin expression. (**M**)Table showing the top 8 hits of ATF occupancy using CUT&RUN sequencing, senescence-associated genes (KEGG map04218) marked in red. Tss=transcription start site; dist=distance;. (**N, P**) Enrichment plots of senescence genes and (**O, Q**) corresponding heat maps (using SenMayo data set) by comparing between siNTC unstimulated vs IL-1a stimulated groups (**N**) and between IL-1a stimulated but either siNTC or siATF4 groups (**P**). (**O**) Heatmap of gene expression levels of genes used in (N). (**Q**) Heatmap of gene expression levels of genes used in (P). In all graphs bars show mean ±SEM; dots show biological replicates or individual experiments. Statistics: Unpaired t-test (I-L) and one way ANOVA, Tukey’s or Holm-Šídák’s multiple comparison test (A-D) ***p>0.001; **p>0.01, *>0.05.

So far, we have observed that the stimulation of NSC with IL-1α has initiated molecular changes that collectively point toward a senescent phenotype [39]. These changes include the metabolic shift towards glycolysis, the expression of several inflammatory genes and the inhibition of proliferation supported by an increase expression of CDK1A/p2^Cip1^. We further interrogated the expression of genes that had been reported to be associated with this senescent phenotype including *Cxcl1*, *Ccl2,* [40] and *Cdkn2a/p16* (Figure 6J-L) [39]. We could determine that a significant increase in expression of all these genes followed a 24h stimulation with IL-1α and could be impaired with ATF4 knockdown.

Using CUT&RUN-sequencing to explore genomic occupancy of ATF4 in IL-1α-stimulated NSCs versus unstimulated controls, we found that two out of top eight genes with the highest TSS occupancy were associated with senescence (KEGG, map04218) (Figure 6M, red text). This further strengthened our hypothesis that IL-1α induces senescence in NSCs via ATF4. To further validate this hypothesis, we performed GSEA analysis on our RNAseq data using the SenMayo senescence gene set [41] (Figure 6N-Q). We found a strong enrichment of senescence genes in IL-1α stimulated NSC versus unstimulated control (NES=2.5, Figure 6N, 6O) and this enrichment was abrogated in IL-1α stimulated NSC with ATF4 knockdown (NES=-2.46, Figure 6P, 6Q).

## Discussion

The hallmarks of cellular senescence involve metabolic changes, cell cycle arrest and release of the senescent associated secretory phenotype (SASP) [42]. Senescence has various triggers, the most common being aging, but also known to occur following inflammatory insults [39]. In the current study, we reveal that IL-1α, at levels detected in rat TBI, induces a senescent phenotype in adult NSCs mediated by the activation of the ISR/ATF4 pathway. Interestingly, the connection between ISR and senescence was discussed in a recently published review[43]. IL-1α, besides being a proinflammatory cytokine, is vastly different from IL-1β, in it being an alarmin, also present in non-hematopoietic cells, and has the capability to induce sterile inflammation [44], crucial for the pathology of TBI. It has gained quite recent attention due to its contribution to the development of neurotoxic A1-type astrocytes [45]. IL-1α induces senescence in various systems and cell types [46] but so far, it has not been associated to the ER or ISR stress in NSCs. However, IL-1α has been demonstrated to induce hepatocyte cell death via activation of Chop [47] and more recently, Almog T et al [48] showed a similar IL-1α-dependent ATF4/Chop driven induction of apoptosis in macrophages, findings that support our results on IL-1α-induced ISR/ATF4 pathway.

Another hallmark of senescence is cell cycle arrest. We observed that in our experiments, IL-1α led to a decrease in NSC proliferation, also corroborated by an increase of the cyclin dependent kinase inhibitor, CDKN1A/p21^Cip1^. The CDKN1A/p21^Cip1^ is a target gene of ATF4 [49] supporting the IL-1α-ATF4-p21 axis we observed in the NSCs. Moreover, OLIG2, a transcription factor regulating self-renewal [50] is suppressed by IL-1α in our experiments. Interestingly, Ligon KL et al [51] demonstrated that OLIG2 suppresses CDKN1A/p21^Cip1^, again in line with our data, since a lowering of OLIG2 levels would promote an icreased expression of Cdkn1a/p21. In conclusion, IL-1α induces ATF4 translation which in turn downregulates OLIG2 expression and upregulates CDKN1A/p21^Cip1^ and leads to decrease of NSC proliferation. Cell proliferation and metabolic activity go hand in hand, and in our experiments, IL-1α-reduced proliferation is in line with the observed glycolytic shift, characteristic for less proliferative NSCs [52].

Transcriptomic analysis on TBI-isolated NSCs reveal an enrichment of inflammation-associated pathways, including TNF. Surprisingly, this occurs at day 28 post-TBI and not much at day 7, when the inflammatory status reaches its peak according to the CSF levels of IL1α and other factors (Figure 2A and data not shown). However, it is well established that in TBI, after the initial inflammatory peak at days 7-14, cellular death and release of alarmins, likely including IL-1α, perpetuate in the penumbra surrounding the contusion injury. In our experiments, this is exemplified by a massive cortical lesion at day 28, (Supplementary Figure 2I). This suggests that the NSCs have been most likely exposed to local inflammation for 21 days, even though the inflammation is not measurable in the CSF. This could explain why NSCs display the more dramatic transcriptomic changes at the late time-point rather than at day 7 post-TBI.

Finally, the signal transduction from IL-1α leading to ATF4 translation is mediated, at least in part, via the eIF2α enzyme, reversed by the small molecule ISRIB, an ISR inhibitor [53]. We had explored the activation of the mammalian target of rapamycin (mTOR) pathway since IL-1α can trigger this pathway [54] and additionally, ATF4 translation can be induced via mTOR (mTORC1) [55]. Even if our results were inconclusive, we cannot exclude that mTOR was activated in NSCs following IL-1α. Interestingly, studies on human neural progenitor cells revealed that these cells expressed senescence markers in primary progressive MS and in culture *in vitro*. If treated with rapamycin, an mTOR inhibitor, the senescent phenotype was reversed [56] which ties back to the ATF4 as being a senescence trigger.

In the current study, we have evidence for a new link between IL-1α/ISR/ATF4 orchestrating the senescent phenotype in NSCs during TBI-related neuroinflammation. This system has high potential for therapeutic exploration, since there are Food and Drug Administration (FDA) approved drugs available both for the IL-1 system (Anakinra, IL-1 receptor antagonist) and ISR (Trazodone).

## Acknowledgements

We would like to thank Guangbei Cheng for excellent cellular work, Britt Meyer and Paula Mannström for tissue sectioning, Annika van Vollenhoven for FACSorting, the staff at the animal house facilities, AKM at Karolinska University Hospital and KMB at Karolinska Institutet.

