## Supplementary figures for "ATF4 orchestrates IL-1α-induced senescence in adult neural stem cells"

### *Supplementary figure 1.*

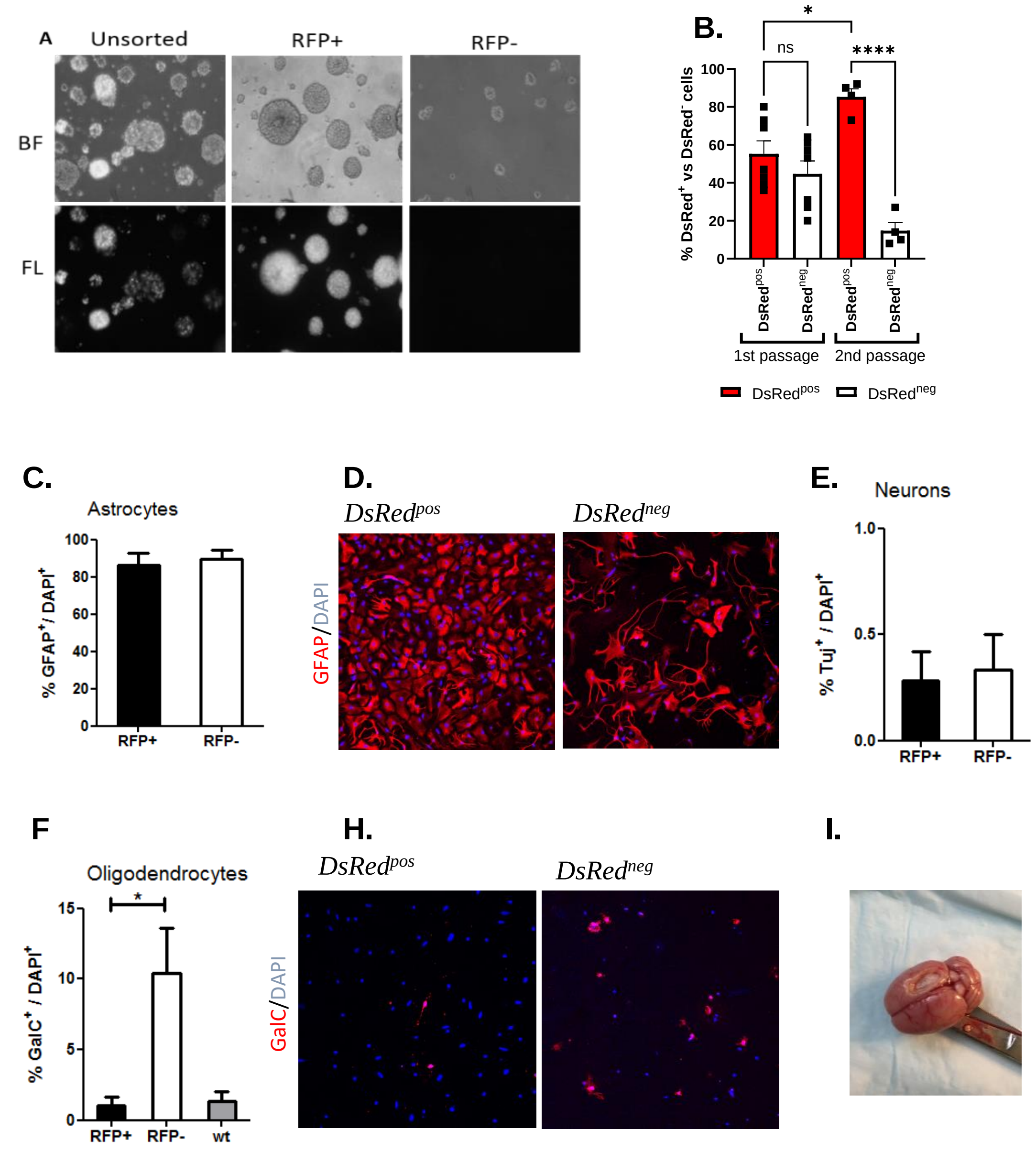


***Supplemtary figure 1. RedSox NSC culture characterization** (**A**). Representative NSCs cultures from the RedSox rat. BF=bright field; FL=fluorescence. (**B**).Percentage of DsRed$^{pos}$ vs DsRed$^{neg}$ NSC cultures following the first and second culture passage. N=4-6 cultures. Differentiation of RedSox NSC sorted into DsRed$^{pos}$ or DsRed$^{neg}$ cells or NSCs from wild type rats. The cells were kept in differentiation media 5-7 days following the second culture passage (**C**) Graph showing percentage of GFAP+ of total number of cells (DAPÍ+) cells. (**D**) Representative pictures of GFAP immunolabeled astrocytes in red, DAPI labeled nuclei in blue. (**E**) Percentage of Tuj+ neurons in differentiated NSC populations DsRed$^{pos}$ (RFP+) and DsRed$^{neg}$ (RFP-). (**F**) Percentage of GalC+ oligodendrocytes out of total number of cells (DAPI+) (H) Representative pictures of differentiated NSC immunoleabeled for GalC (red) oligodendrocytes and DAPI (blue) nuclei. Unit bar= (I) Picture showing the brain of a TBI rat at day 28 post-injury.*

#### *Supplementary figure 2.*

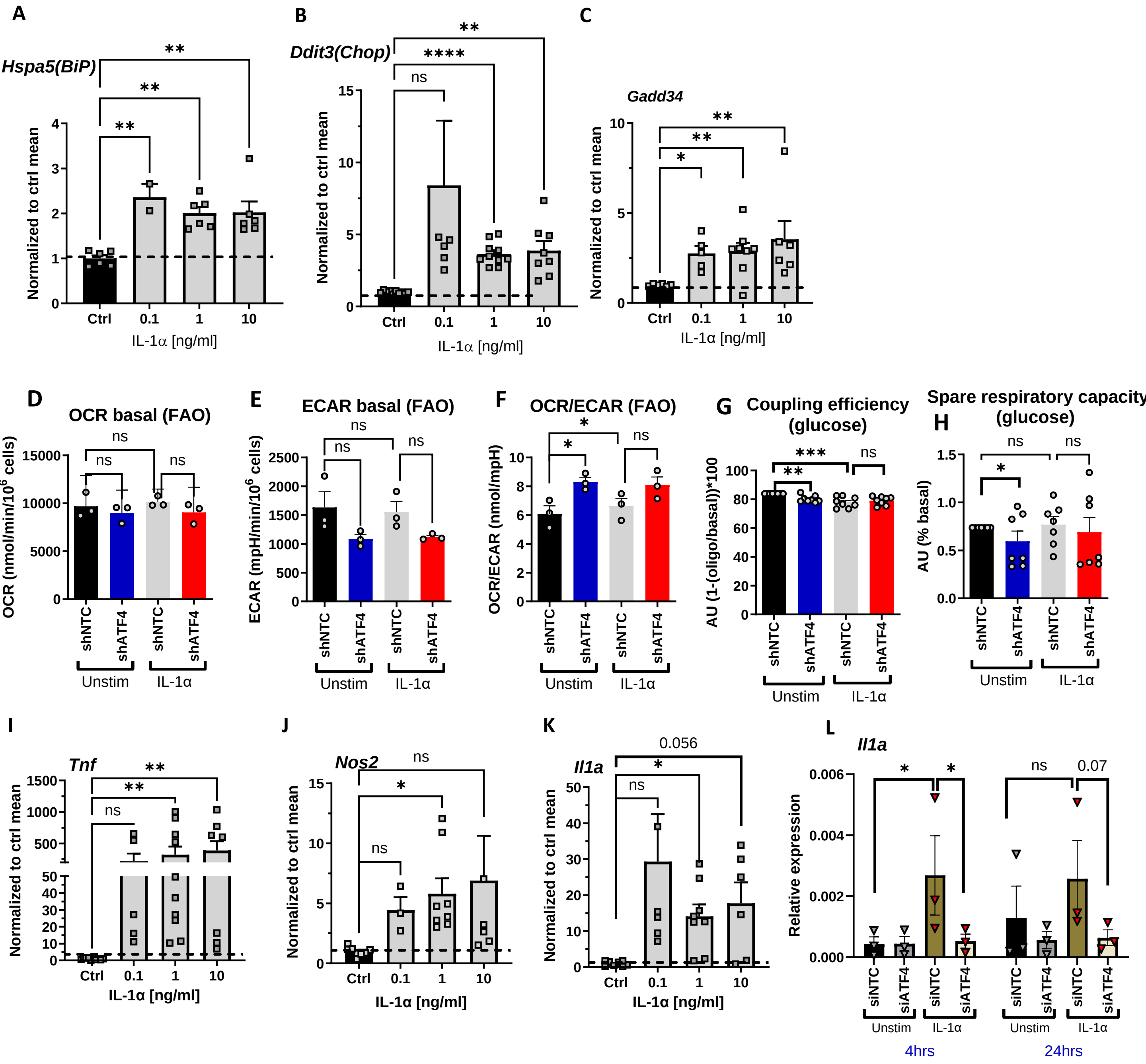


***Supplementary figure 2.*** *Graphs show the RNA expression level measured with real-time RT-PCR and normalized to βactin following a 24h stimulation with increasing concentrations of IL1a.* ***(A)*** *Hspa5(BiP),* ***(B)*** *Ddit3(Chop),* ***(C)****Gadd34,* ***(I)*** *Tnf,* ***(J)*** *Nos2* ***(K)*** *Il1a.* ***(L)*** *Real-time PCR analysis of IL1a expression in NSC that have been stimulated with IL-1a or left unstimulated after transfection with siNTC or siATF4. In all graphs bars show mean ±SEM; dots show biological replicates or individual experiments. Statistics: One way ANOVA, Tukey's or Holm-Šídák's multiple comparison test ***p>0.001; **p>0.01, *>0.05.*

*(**D-F**). Fatty acid oxidation (FAO) assessment using palmitate as energy source. Basal OCR (**D**), ECAR (**E**) and ratio between OCR and ECAR in NSCs transduced with shATF4 or shNTC plasmids, stimulated with IL-1a or kept unstimulated (F). (**G;H**) Glucose metabolism assessment. (**D**) Coupling efficiency and (**E**) spare respiratory levels. AU=area under curve; oligo=oligomycin. In all graphs bars show mean ±SEM; dots show biological replicates or individual experiments. Statistics: One way ANOVA, Tukey's or Holm-Šídák's multiple comparison test ***p>0.001; **p>0.01, *>0.05.*
